# Dichotomic Regulation of Striatal Plasticity by Dynorphin

**DOI:** 10.1101/2022.04.04.487051

**Authors:** Renzhi Yang, Rupa R. Lalchandani Tuan, Fuu-Jiun Hwang, Daniel W. Bloodgood, Dong Kong, Jun B. Ding

## Abstract

Modulation of corticostriatal plasticity alters the information flow throughout basal ganglia circuits and represents a fundamental mechanism for motor learning, action selection, and reward. Synaptic plasticity in the striatal direct- and indirect-pathway spiny projection neurons (dSPNs and iSPNs) are dichotomically regulated by two distinct networks of GPCR signaling cascades. While it is well-known that dopamine D2 and adenosine A2a receptors bidirectionally regulate iSPN plasticity, it remains unclear how D1 signaling modulation of synaptic plasticity is counteracted by a dSPN-specific Gi signaling. Here, we show that striatal dynorphin selectively suppresses long-term potentiation (LTP) through Kappa Opioid Receptor (KOR) in dSPNs. Both KOR antagonism and conditional deletion of dynorphin in dSPNs enhance LTP counterbalancing with different levels of D1 receptor activation. Behaviorally, mice lacking dynorphin specifically in dSPNs show normal motor behavior and reward-based learning, but enhanced flexibility during reversal learning. These findings support a model in which D1R and KOR signaling bidirectionally modulate synaptic plasticity in striatal direct pathways and behavior.

## INTRODUCTION

The striatum is the gateway into the basal ganglia and receives convergent glutamatergic inputs from the cortex and thalamus. Activity-dependent excitatory synaptic plasticity in the striatum alters the information transfer through the basal ganglia and is crucial for fine motor control, motor learning, habit formation, and action selection (Graybiel and Grafton, 2015; Gremel and Costa, 2013; Kauer and Malenka, 2007). Maladaptive and/or loss of synaptic plasticity are thought to mediate motor symptoms seen in movement disorders, such as Parkinson’s disease and L-DOPA-induced dyskinesia (Giordano et al., 2018; Girasole et al., 2018; Pisani et al., 2005; Surmeier et al., 2014). In the striatum, more than 90% of neurons are spiny projection neurons (SPNs), forming two parallel pathways characterized by their axonal projections, expression of neuropeptide and neuromodulatory receptors (Gerfen and Surmeier, 2011). In particular, dynorphin and dopamine D1 receptor are exclusively expressed in direct-pathway SPNs (dSPNs), whereas enkephalin, dopamine D2 receptor, and adenosine A2a receptor are exclusively expressed in indirect-pathway SPNs (iSPNs) (Zhai et al., 2019). This pathway-specific expression of neuromodulatory receptors is optimally positioned to regulate the excitatory synaptic plasticity (Gerfen and Young, 1988). For example, in iSPNs, activation of Gs-coupled adenosine A2a receptor promotes long-term potentiation (LTP) expression, and activation of Gi-coupled D2 receptor suppresses LTP expression (Higley and Sabatini, 2010; Kozorovitskiy et al., 2015; Shen et al., 2008; Surmeier et al., 2014). Conversely, what remains unclear is whether there is a dSPN-specific counterpart of D2R that suppresses LTP expression. One candidate is the Gi-coupled M4 muscarinic receptor (M4R), which has been shown to successfully shunt LTP in dSPNs (Shen et al., 2015). The effect of M4R is assumed to be low in iSPNs because it is preferentially expressed in dSPNs (Yan et al., 2001; Zhai et al., 2019). However, although M4R show highly differential expression levels in dSPN and iSPNs, M4R is also expressed in iSPNs, besides its extensive expression at corticostriatal axonal terminals and striatal cholinergic interneurons (Bernard et al., 1992; Hersch et al., 1994), which makes M4R ill-fitted for direct-pathway specific modulation.

Another pathway-specific candidate is the Gi-coupled Kappa Opioid Receptor (KOR). KOR is also preferentially expressed in dSPNs (Oude Ophuis et al., 2014; Tejeda et al., 2017), and importantly, the endogenous ligand, dynorphin (Dyn), is exclusively expressed in dSPNs (Reiner and Anderson, 1990). Even though the distinct expression pattern of Dyn in the dSPNs is well-known, the endogenous function of Dyn in the striatal direct pathway is still not well understood. Previous studies largely focused on its acute regulation of synaptic transmission and synaptic plasticity. For example, activation of KOR by Dyn can suppress dopamine (DA) release in the striatum (Di Chiara and Imperato, 1988; Spanagel et al., 1992) and the effect is short-term and can be reverted after washout (Ehrich et al., 2015). In addition, acute KOR activation can also suppress glutamatergic synaptic transmission (Coleman et al., 2021; Mu et al., 2011) or prolonged activation of KOR causes long-term depression (LTD) (Atwood et al., 2014) through pre-synaptic mechanisms.

Here, we used a combination of pharmacological manipulation of KOR and a striatal direct-pathway specific Dyn conditional knockout (cKO) mouse to probe the endogenous function of Dyn, in particular, we focus on excitatory synaptic transmission and plasticity. Interestingly, we found that indeed M4R activation can also suppress spike-timing-dependent LTP (STDP-LTP) in iSPNs, suggesting that M4R mediated suppression of LTP is not pathway-specific. In contrast, we found that either blockade of KOR receptors, or direct-pathway-specific deletion of Dyn can selectively suppress STDP-LTP in dSPNs, but not in iSPNs. Furthermore, we characterized the interaction of Dyn/KOR with D1R signaling in dSPNs and found that Dyn/KOR counterbalances with different levels of D1R activation and result in different levels of STDP-LTP expression. Finally, we found mice lacking Dyn specifically in dSPNs (Dyn cKO mice) showed normal motor behavior, working memory, anxiety level, and normal reward-based learning. But interestingly Dyn cKO mice showed enhanced performance in re-learning a reversed reward contingency. Taken together, our studies provide novel mechanistic insights into dichotomic regulation of striatal direct-pathway synaptic plasticity by Dyn/KOR signaling and demonstrate a link between endogenous function of striatal Dyn, striatal direct pathway synaptic plasticity and behavioral cognitive flexibility.

## RESULTS

To induce LTP in the dorsal striatum, we utilized spike-timing-dependent plasticity (STDP) protocol combined with perforated-membrane patch clamp recording (Shen et al., 2008). The induction protocol comprised of a series of 50 Hz bursts of “pre-post” pairing, in which three presynaptic inputs preceded three postsynaptic spiking by 5 ms (Figure 1A). Presynaptic glutamatergic afferent fibers were stimulated with a small theta glass pipette close to the recorded SPN (50-100 µm), and postsynaptic spiking was induced by current injection through the perforated patch. Before induction, we recorded theta glass-evoked excitatory postsynaptic currents (EPSCs) to obtain a stable 5-minute baseline, and after STDP-LTP induction, we continued to record the evoked EPSCs for at least 30 minutes to monitor changes in EPSC amplitudes. To distinguish dSPNs and iSPNs, we used D1-Cre;Ai32 mice, in which Channelrhodopsin-2 (ChR2) is specifically expressed in D1 neurons, enabling the identification of dSPNs and iSPNs based on the presence or absence of the optically evoked response, respectively (please see Methods for details). In both dSPNs and iSPNs, this STDP protocol reliably induced LTP (Figure 1B,F).

**Figure 1.**
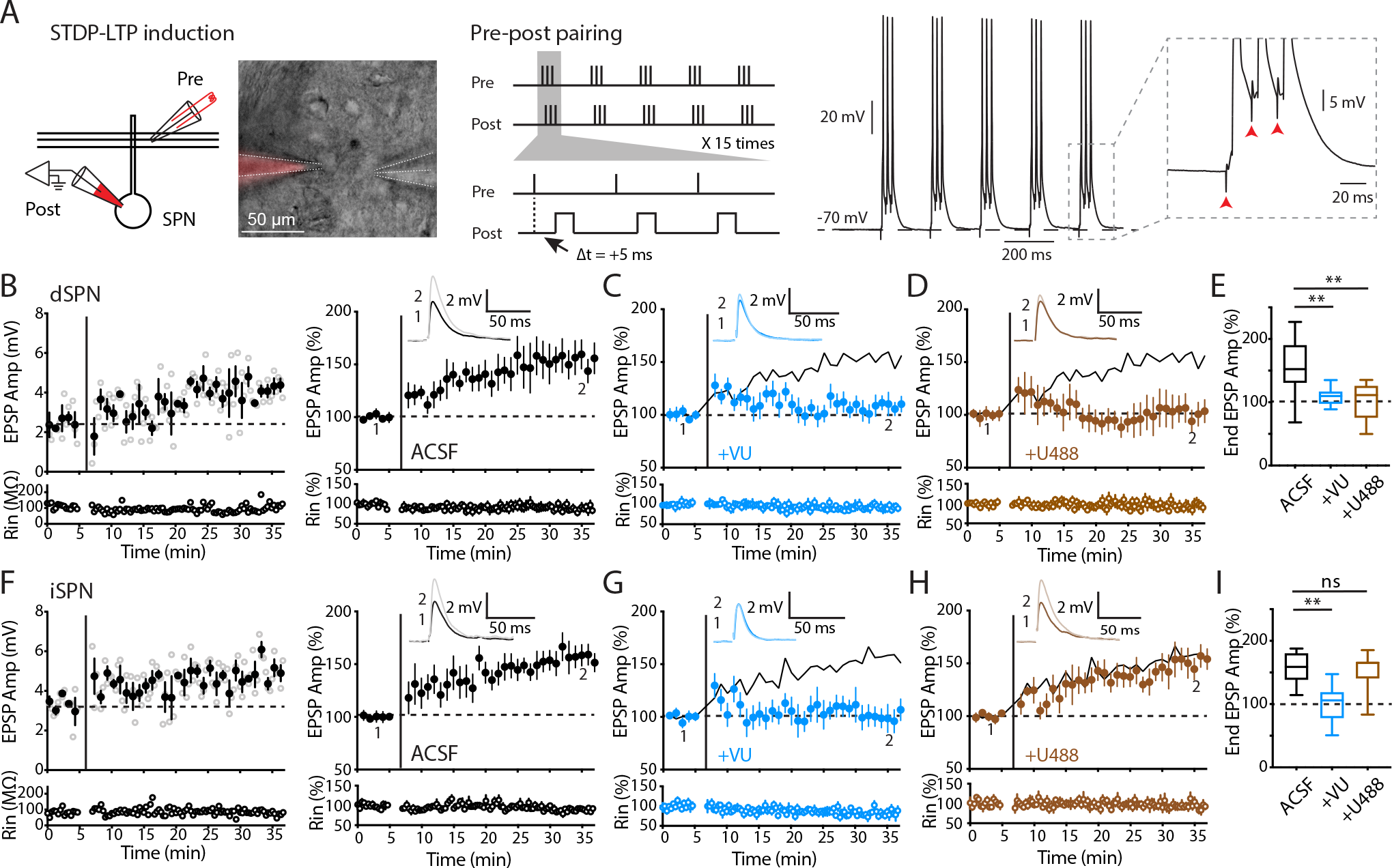
Pathway-specific modulation of striatal STDP-LTP by KOR. (**A**) Left: the schematic and image depicting perforated patch recording from a postsynaptic SPN with presynaptic stimulation using a two-barrel theta glass. Middle: the theta-burst spike-timing-dependent plasticity (STDP) pairing protocol for LTP induction. Right: a representative trace showing action potentials and EPSPs recorded during STDP-LTP induction. (**B, F**) Left: example experiments of dSPN (B) and iSPN (F) recordings from slices obtained from WT mice. Open circles show EPSP amplitudes (Amp) and input resistance (Rin) throughout the recording. Filled circles are averages of 3 trials (± SEM). STDP-LTP induction is indicated by the vertical bar. The dashed line is the average baseline EPSP amplitude before induction. Right: summary plots of the normalized EPSP amplitudes and Rin as a function of time. Inset: representative average EPSP traces of the baseline (1) and the last 5 min (2). (**C, G**) Summary plots of recordings obtained from WT mice with administration of M4R PAM (VU10010). VU10010 suppressed LTP in both dSPNs (C) and iSPNs (G). Solid lines are control LTP for reference. (**D, H**) Summary plots of recordings obtained from WT mice with administration of KOR agonist (U50488). U50488 suppressed LTP in dSPNs (D) but not iSPNs (H). Solid lines are control LTP for reference. (**E, I**) Box-plot summaries showing the normalized EPSP amplitudes from the last 5 min of recording (end EPSP). (E) dSPN LTP was suppressed by VU10010 and U50488. Control (ACSF), n = 12/8, 153.04% ± 12.64; +VU, n = 10/4, 109.37% ± 4.42; +U488, n = 8/4, 102.09% ± 10.59. Control vs +VU, p = 0.0034; control vs +U488, p = 0.0055, Mann-Whitney. (I) iSPN LTP was suppressed by VU10010 but not U50488. Control (ACSF), n = 8/5, 156.43% ± 8.76; +VU, n = 9/4, 99.98% ± 9.77; +U488, n = 9/5, 151.99% ± 9.79. Control vs +VU, p = 0.001; control vs +U488, p = 0.96, Mann-Whitney. Data are presented as mean ± SEM. ns: not significant, p > 0.05, ** p < 0.01.

### KOR suppresses LTP specifically in dSPNs

To investigate whether the activation of Gi-coupled M4R can differentially regulate LTP in a pathway-specific manner, we incubated the brain slices obtained from wildtype (WT) mice with the M4R positive allosteric modulator (PAM) VU10010 (VU, 5 µM), which enhances the response to endogenous acetylcholine. Consistent with the previous finding (Shen et al., 2015), the M4R PAM VU10010 blunted LTP expression in dSPNs (Figure 1C,E). It has been generally assumed that the effect of M4R is small in iSPNs because M4R is preferentially expressed in dSPNs (Yan et al., 2001; Zhai et al., 2019). However, on the contrary, we observed a similar level of suppression of LTP in iSPNs by M4R PAM VU10010 (Figure 1G,I). This result indicated that M4R activation broadly regulates LTP in both dSPNs and iSPNs.

Because of the unique expression pattern of Dyn (Reiner and Anderson, 1990) and preferential expression of KOR in the dSPNs (Oude Ophuis et al., 2014; Tejeda et al., 2017), we next tested the effect of KOR activation on LTP in both dSPNs and iSPNs in WT mice. We used the same recording condition and replaced M4R PAM with KOR agonist U50488 (U488, 1 µM). We observed that U488 abolished LTP expression in dSPNs (Figure 1D,E), while LTP in iSPNs remained intact (Figure 1H,I), indicating that KOR signaling regulates striatal LTP in a pathway-specific manner.

### D1-pDyn cKO enhances LTP specifically in dSPNs

One caveat associated with KOR agonists is that KOR activation can directly affect the basal glutamatergic and dopaminergic synaptic transmission (Atwood et al., 2014; Mu et al., 2011; Spanagel et al., 1992). To confirm the pathway-specific regulation of LTP by endogenous KOR signaling in the striatum, we next used genetic tools to selectively remove the endogenous KOR ligand, Dyn, by generating a striatum direct-pathway-specific Dyn conditional knockout mice.

The dynorphin family consists of several variants derived from the prodynorphin (pDyn) gene (Healy and Meador-Woodruff, 1994), all of which share the amino acid sequence encoded in the exon 4, conferring them the ligand binding specificity to KORs (Schwarzer, 2009). Therefore, we generated pDyn-floxed (pDyn^fl/fl^) mice by flanking exon 4 of the pDyn gene with loxP sites (Figure 2A). We specifically knocked out Dyn in the striatum by crossing the pDyn^fl/fl^ mice with D1-Cre mice, which express Cre recombinase in D1 dopamine receptor (D1R)-containing cells. While D1R is also expressed at low levels in several other brain areas (Gong et al., 2007), dynorphin and D1R co-expression occur mainly in the striatal dSPNs, thus crossing the pDyn^fl/fl^ mice with D1-Cre mice generated D1-pDyn cKO mice that lacks dynorphin specifically in the striatal direct pathway (Figure 2B).

**Figure 2.**
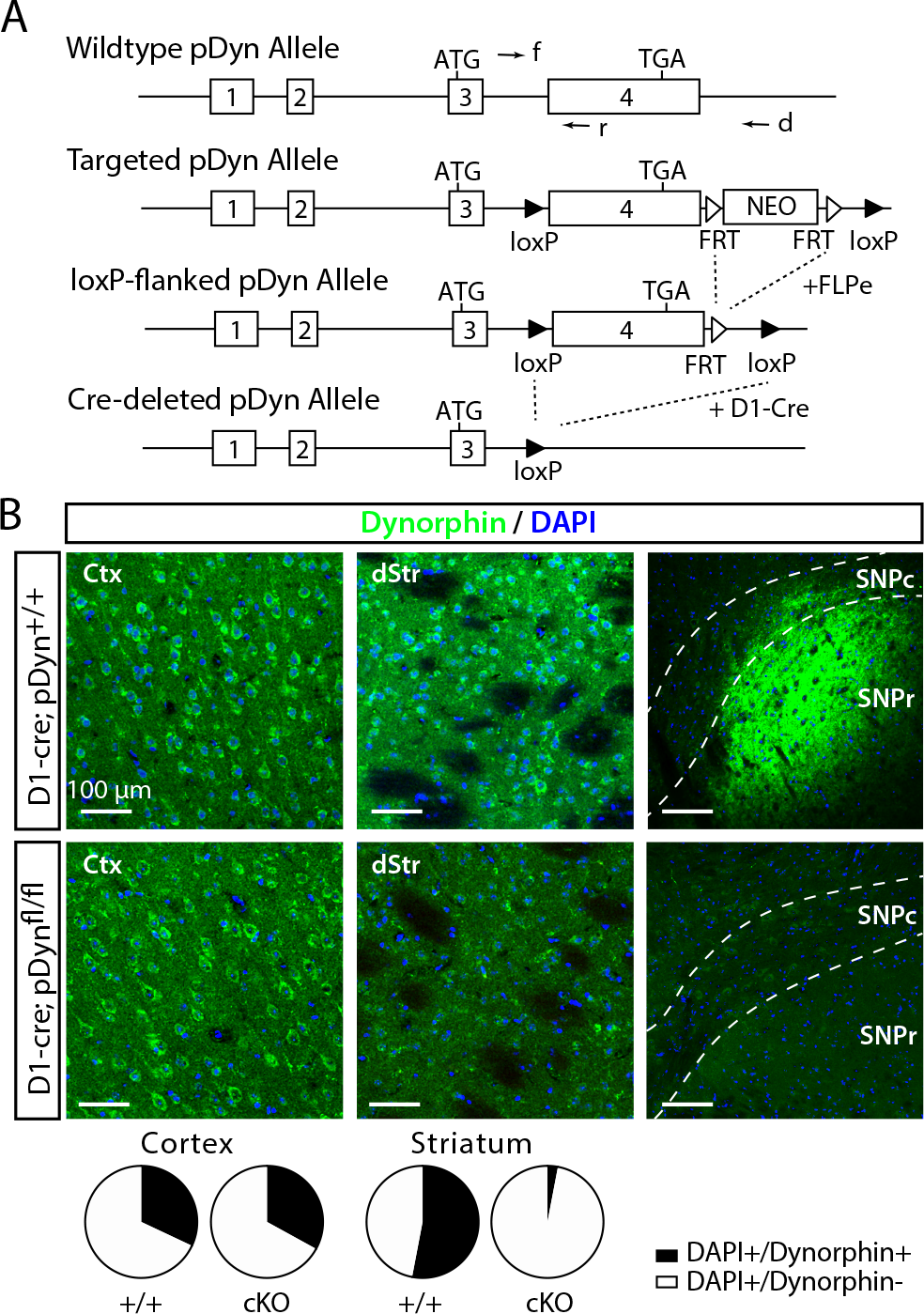
Direct pathway-specific deletion of pDyn. (**A**) Gene targeting strategy scheme for pDyn conditional knockout (cKO) mice, including the diagrams of wildtype pDyn allele, targeted allele, loxP-flanked allele, and Cre-deleted allele. In the targeted allele, pDyn Exon 4 and a neo cassette are flanked by loxP sites. The neo cassette is removed from the targeted allele by FLP recombinase. pDyn exon 4 is then deleted in D1 cells by breeding to a D1-Cre mouse line. The primers used for genotyping are also indicated as arrows. (**B**) Immunohistochemistry of dynorphin in striatum and cortex. Top: sample images of a coronal section of cortex (left), striatum (middle) and substantia nigra (right), stained for dynorphin (green) and DAPI (blue). Bottom: quantification of co-labelling of dynorphin with DAPI in the cortex (WT, n = 3500 cells / 4 mice, 31.6%; cKO, n = 3500 cells / 4 mice, 33%) and striatum (WT, n = 4002 cells / 4 mice, 52.9%; cKO, n = 3996 cells / 4 mice, 2.9%).

To assess the deletion efficiency in the striatum, we performed immunostaining with anti-Dyn antisera. In WT control striatal sections, approximately half of the DAPI labelled neurons were dynorphin positive (WT control: 52.9%; Figure 2B), which is consistent with previous reports (Gerfen and Surmeier, 2011). By contrast, Dyn expression in the striatum of D1-Cre;pDyn^fl/fl^ mice was almost fully abolished in the striatum (cKO: 2.9%; Figure 2B). We also immunostained and imaged the substantia nigra region, where the axonal projections of dSPNs terminate, and again observed a diminished Dyn signal in cKO slices. Cortical dynorphin expression on the other hand was not affected in the D1-Cre;pDyn^fl/fl^ mice (WT: 31.6%, cKO: 33%; Figure 2B), confirming that deletion of Dyn in D1-Cre;pDyn^fl/fl^ mice was indeed specific to the striatum.

We recorded STDP-LTP in dSPN from the D1-Cre;pDyn^fl/fl^ mice. Strikingly, we found that LTP in dSPNs was significantly enhanced in the Dyn cKO mice (Figure 3A,B). We also pharmacologically blocked KOR with the antagonist norbinaltorphimine (NorBNI, 1 µM) in the D1-Cre;pDyn^+/+^ mice, and found similar effect, norBNI enhanced LTP in dSPNs (Figure 3C,D). These results are complementary to the result above with KOR activation (Figure 1D,E). Together, these data show that LTP is bi-directionally regulated by KOR signaling: Dyn/KOR signaling suppresses LTP expression in dSPNs and pharmacologically blocking KOR or genetically removing its ligand Dyn enhances LTP.

**Figure 3.**
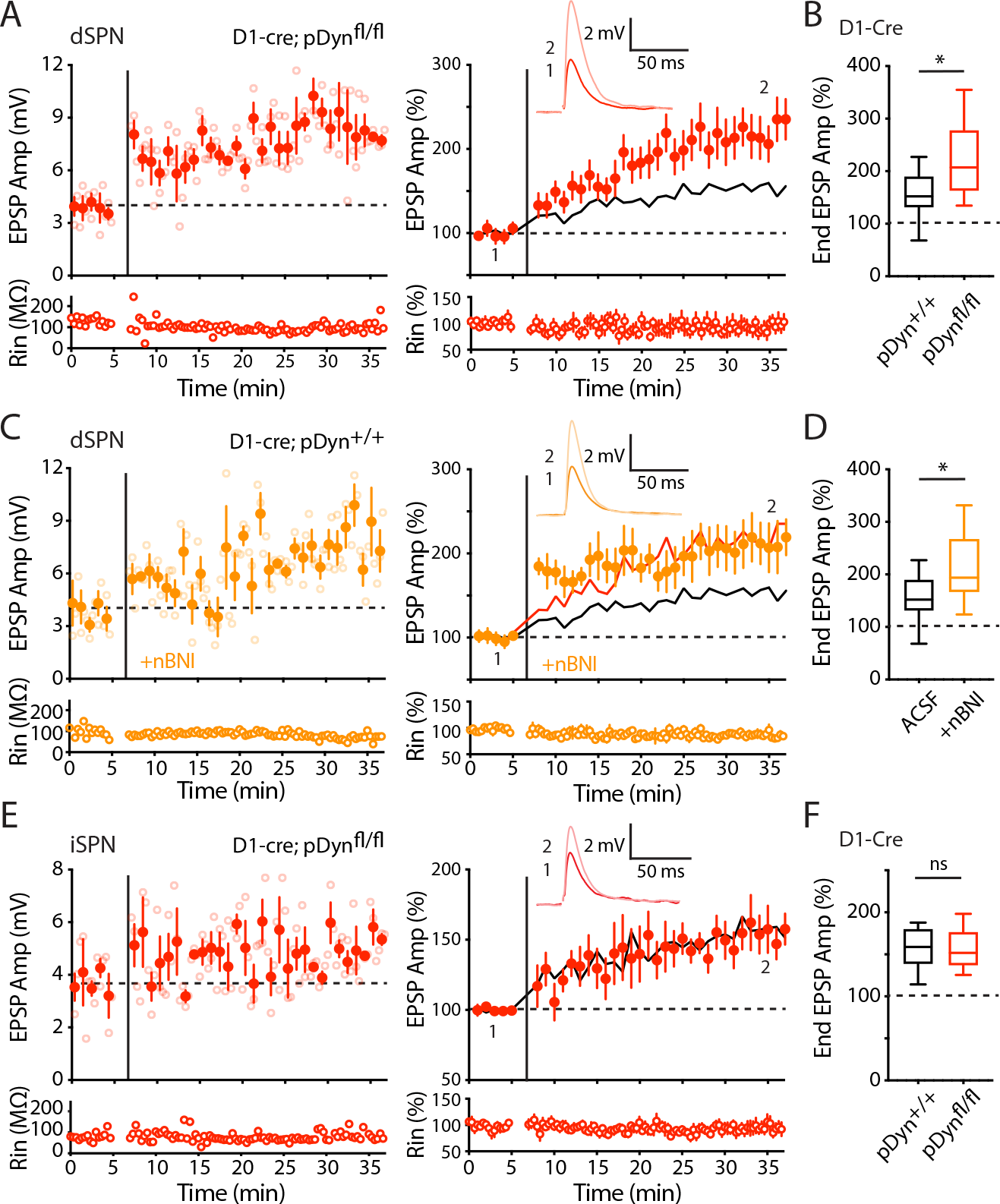
D1-pDyn cKO enhanced LTP in dSPNs, but not iSPNs. (**A**) Left: an example experiment of dSPN recordings from slices from pDyn cKO mice (D1-Cre;pDyn^fl/fl^). Right: summary plot showing an enhanced LTP in cKO compared with WT (D1-Cre;pDyn^+/+^)group. The solid line shows the control LTP from Figure 1(B) for reference. (**B**) Box-plot summary showing enhanced dSPN LTP by D1-pDyn cKO. WT, n = 12/8, 153.04% ± 12.64; cKO, n = 10/7, 220.86% ± 23.28. p = 0.0206, Mann-Whitney. (**C**) Left: an example experiment of dSPN recordings from WT mice (D1-Cre;pDyn^+/+^) with administration of KOR antagonist (+nBNI). Right: summary plot showing that NorBNI significantly enhanced the LTP to a similar level as cKO. The solid black and red lines are LTP recorded from WT and cKO mice, respectively, for reference. (**D**) Box-plot summary showing enhanced dSPN LTP by norBNI. Control (ACSF), n = 12/8, 153.04% ± 12.64; +nBNI, n = 8/3, 212.98% ± 23.66. control vs +nBNI, p = 0.0473, Mann-Whitney. (**E**) Left: an example experiment of iSPN recordings from cKO slices. Right: summary plot showing similar LTP levels in control and cKO. The solid line is the control LTP from Figure 1F for reference. (**F**) Box-plot summary showing similar iSPN LTP between control and cKO. WT, n = 8/5, 156.43% ± 8.76; cKO, n = 8/6, 156.84% ± 8.81. WT vs cKO, p = 0.88, Mann-Whitney. Data are presented as mean ± SEM. ns: not significant, p > 0.05, * p < 0.05.

In addition, we recorded LTP in iSPNs from the D1-Cre;pDyn^fl/fl^ mice. LTP in iSPNs of cKO mice were comparable to WT control mice (Figure 3E,F), confirming that suppression of LTP by dyn/KOR signaling is pathway specific.

### No change on basal dopaminergic and glutamatergic transmission in D1-Cre;pDyn^fl/fl^ mice

Previous studies have shown that acute activation of dyn/KOR signaling can suppress the dopamine (DA) (Di Chiara and Imperato, 1988; Schlosser et al., 1995; Spanagel et al., 1992) and glutamatergic release (Atwood et al., 2014; Hjelmstad and Fields, 2001, 2003; Mu et al., 2011). To test whether dopaminergic and glutamatergic signaling are affected in D1-pDyn cKO mice, we measured basal dopaminergic and glutamatergic transmission. We first used fast scan cyclic voltammetry (FSCV) to assess dopaminergic transmission in the dorsal striatum. We used a local bipolar electrode to evoke DA release with a single stimulation or a train of three stimulations at 50 Hz and measured the evoked local extracellular DA concentration ([DA]o). We did not find any differences in evoked [DA]o between WT and cKO mice (Figure S1A-C,E). We also did not observe any differences in paired-pulse ratio (PPR) (Figure S1D). We next recorded the spontaneous miniature excitatory postsynaptic currents (mEPSCs) (Figure S2A,B) to assess basal glutamatergic transmission. We did not find any significant differences in mESPC mean amplitude or frequency in either SPN types between WT control and cKO mice (Figure S2C,D). Furthermore, using 2-photon imaging in acute brain slices in which SPNs were filled with the red fluorophore Alexa594 through the whole-cell recording pipette, we examined the density of spiny protrusions along SPN dendrites, where most glutamatergic synapses are formed on SPNs. We found that there was no significant differences in spine density in either dSPNs or iSPNs between WT and cKO mice (Figure S2E,F). These results indicated that deletion of striatal Dyn did not affect the basal levels of dopaminergic or glutamatergic transmission in the dorsal striatum. Our results thus provide evidence that the enhanced LTP expression in dSPNs in D1-pDyn cKO mice is not due to an altered baseline DA or glutamatergic release, but rather suggests a direct effect of autonomous activation of KOR signaling in dSPNs.

### Interaction between Dyn/KOR and D1R signaling on LTP induction

Signaling balance between G-protein coupled receptor (GPCR) cascades controlling striatal synaptic plasticity (Zhai et al., 2019). In dSPNs, D1Rs are crucial for LTP induction (Shen et al., 2008). D1Rs activation promotes LTP induction while D1Rs blockade leads to LTD even with STDP-LTP induction protocols (Shen et al., 2008). In our STDP experiment described above, D1Rs are activated by dopamine released from *en passant* dopaminergic fibers stimulated during STDP induction. However, dopamine levels might be considerably different in physiological conditions. For example, midbrain DA neurons exhibit phasic bursts of firing in response to unexpected rewards and decreased firing rate when an expected reward is omitted (Schultz et al., 1997; Watabe-Uchida et al., 2017), which might result in transient high-level activation or suppression of D1R signaling in dSPNs. Therefore, we investigate how D1R signaling interacts with Dyn/KOR signaling to modulate LTP expression at different D1R activation levels. We treated the striatal slices with the D1R full agonist SKF81297 (SKF, 3 µM) or antagonist SCH23390 (SCH, 3 µM) to examine whether dSPN LTP is modulated by KOR signaling when D1R is fully activated or blocked, respectively. In the presence of the D1R agonist, SKF81297, we observed that both WT control and cKO dSPNs have enhanced levels of LTP (Figure 4A,B) with a similar magnitude (Figure 4D). The finding that the D1R agonist did not further enhance LTP suggests that high levels of D1R activation occlude the LTP enhancement by Dyn deletion. Next, we examine LTP induction and expression when both D1R and KOR are activated by the administration of both agonists SKF 81297 and U50488. Interestingly, U50488 failed to suppress LTP in the presence of the D1R agonist (Figure 4C). Notably, all groups showed comparable high levels of LTP expression regardless of whether KOR is activated or blocked (Figure 4D). Together, these results suggest that D1R signaling dominates the KOR pathway when both are activated and that high-level D1R signaling promotes LTP expression and diminished the suppressive role of Dyn/KOR signaling (Figure 4E).

**Figure 4.**
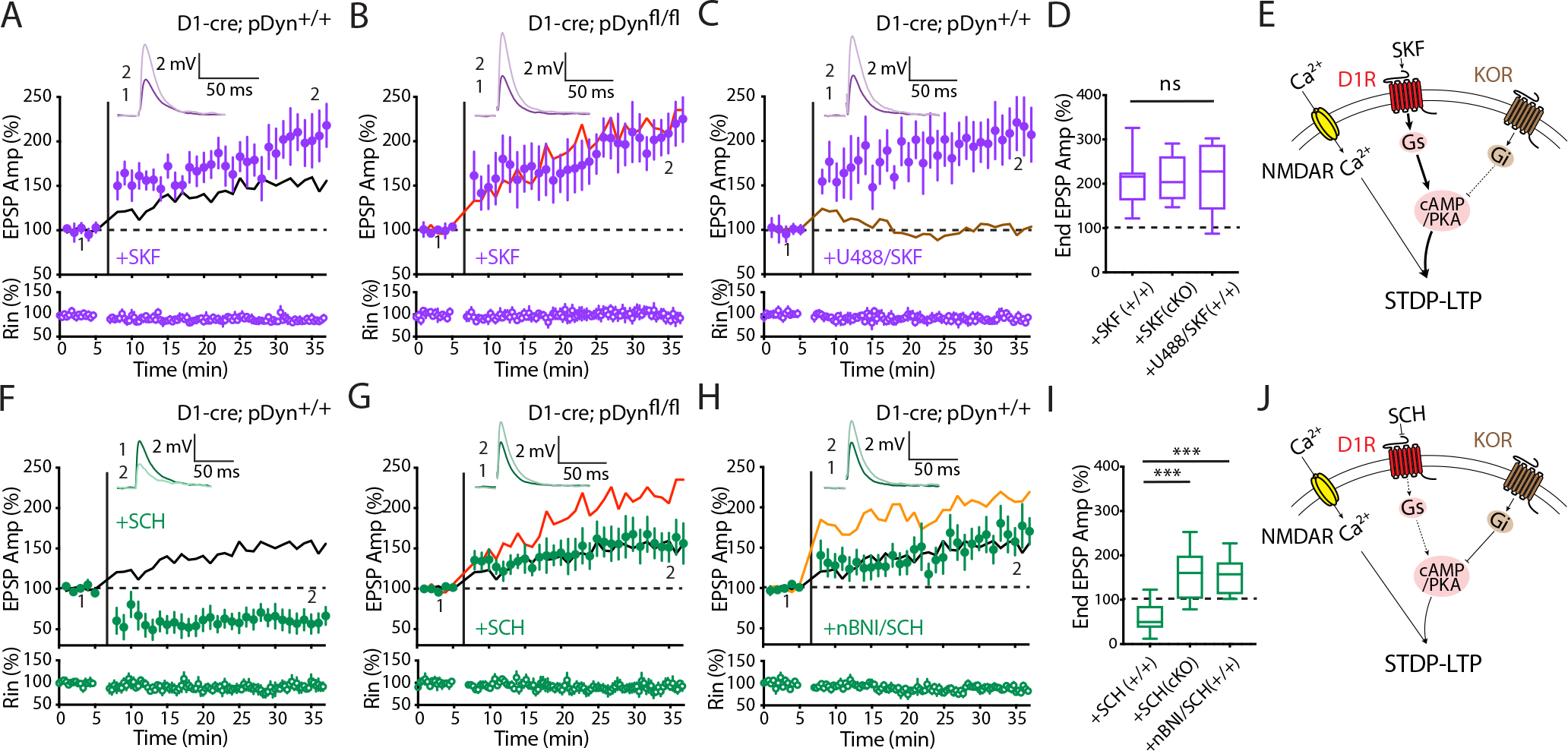
D1R signaling modulated the effect of dyn/KOR on dSPN LTP. (**A**) Summary plot showing enhanced LTP in the presence of D1 agonist (SKF81297) recorded from WT (D1-Cre;pDyn^+/+^)mice. The solid line shows LTP recorded without SKF81297 for reference. (**B**) Summary plot showing enhanced LTP in the presence of D1 agonist (SKF81297) recorded from cKO (D1-Cre;pDyn^fl/fl^)mice. SKF81297 did not further enhance LTP in cKO mice. The solid line shows LTP recorded from cKO mice without SKF81297 for reference. (**C**) Summary plot showing enhanced LTP in the presence of D1 agonist (SKF81297) and KOR antagonist (U50488) recorded from WT (D1-Cre;pDyn^+/+^) mice. In the presence of SKF81297, U50488 failed to block the LTP. The solid line shows recording in the presence of +U50488 alone for reference. (**D**) Box-plot summary showing LTP levels in the presence of SKF81297. WT+SKF, n = 7/3, 206.76% ± 24.5; cKO+SKF, n = 7/3, 211.83% ± 19.76; WT+U488+SKF, n = 7/2, 210.82% ± 29.05. WT+SKF vs cKO+SKF, p = 0.805; WT+SKF vs WT+U488+SKF, p = 0.71; cKO+SKF vs WT+U488+SKF, p = 0.99, Mann-Whitney. (**E**) The schematic depicting the signaling pathway of D1R and KOR activation. D1R activation enhanced dSPN LTP expression regardless of whether dyn/KOR pathway is activated. (**F**) Summary plot showing the same STDP induction protocol resulted in LTD in the presence of D1R antagonist (SCH23390) recorded from WT (D1-Cre;pDyn^+/+^) mice. The solid line shows control LTP for reference. (**G**) Summary plot showing LTP is dampened to WT control LTP level in the presence of D1 antagonist (SCH23390) recorded from cKO (D1-Cre;pDyn^fl/fl^) mice. The solid red line shows LTP in cKO mice and solid black line shows WT control LTP for reference. (**H**) Summary plot showing LTP recorded from WT (D1-Cre;pDyn^+/+^) mice in the presence of both D1 antagonist (SCH23390) and KOR antagonist (norBNI). When D1R is blocked, blocking KOR resulted in normal LTP. The solid orange line shows WT+NorBNI LTP and solid black line shows control LTP for reference. (**I**) Box-plot summary showing LTP levels in the presence of D1R antagonist (SCH23390). Loss of KOR function rescued LTP in the presence of D1R antagonist. WT+SCH, n = 9/4, 59.95% ± 11.72; cKO+SCH, n = 9/3, 156.01% ± 19.31; WT+norBNI+SCH, n = 9/4, 154.68% ± 14.14. WT+SCH vs cKO+SCH, p = 0.0008; WT+SCH vs WT+norBNI+SCH, p = 0.0005; cKO+SCH vs WT+norBNI+SCH, p = 0.99, Mann-Whitney. (**J**) The schematic depicting the signaling pathway of D1R and KOR blockade. Blockade of D1R suppressed dSPN LTP, deletion or blockade of KOR rescued LTP in the presence of D1R antagonist. Data are presented as mean ± SEM. ns: not significant, p > 0.05, *** p < 0.001.

Conversely, in the presence of the D1R antagonist SCH23390 (3 µM), dSPNs from WT D1-Cre;pDyn^+/+^ mice expressed long-term depression (LTD) despite the fact that an LTP induction protocol was used (Figure 4F), which is consistent with previous finding (Shen et al., 2008). Interestingly, D1R antagonist SCH23390 reduced but did not fully block the LTP in dSPNs from D1-Cre;pDyn^fl/fl^ mice (Figure 4G). Consistently, when we blocked KOR with norBNI in dSPNs from WT D1-Cre;pDyn^+/+^ mice, LTP was reduced by SCH23390 compared to LTP in the presence of norBNI alone (Figure 4H), but again LTP was not fully blocked by D1 antagonist SCH23390. Notably, unlike the results above with the D1R agonist, where decreasing or increasing dyn/KOR does not alter the predominant effect of D1R activation, here, LTP induction and expression are comparable to WT control levels when Dyn is conditionally deleted from dSPNs or KOR is blocked (Figure 4G,H,I).

In summary, these results showed that signaling of endogenous striatal dynorphin interacts closely with dopamine D1R signaling to modulate striatal plasticity in direct pathway. Dyn/KOR plays a more prominent role in suppressing LTP in the absence or at lower levels of D1R activation, and its effect may be dampened during higher levels of D1R activation (Figure 4J).

### Enhanced instrumental learning in the reversal phase

Having established the role of striatal dynorphin in regulating synaptic plasticity in direct pathway, we next ask, what is the behavioral relevance? One of the essential functions of the dorsal striatum is to facilitate reward-based instrumental learning, in which mice acquire the association between behavioral actions and reward outcomes (Yin et al., 2008). The dorsal striatum has been shown to encode action-outcome (A-O) contingency, action sequence and habit formation (Balleine and O’Doherty, 2010; Graybiel and Grafton, 2015; Gremel and Costa, 2013). Given that LTP is commonly correlated with learning and memory and an essential role of basal ganglia in the motor control (Albarran et al., 2021; Clem et al., 2008; Rioult-Pedotti et al., 2000), we test whether mice lacking Dyn in dSPNs would have any behavioral deficits using a battery of behavioral tests, including motor control, anxiety, working memory and instrumental learning performance.

The Dyn cKO and their WT control littermates performed equally in a variety of behavioral assays, including the open field task for testing locomotion and anxiety (Figure 5A), balance beam for fine movement control (Figure 5B), Y maze for working memory (Figure 5C), and elevated plus maze for anxiety (Figure 5D). There are no statistically significant differences between WT control (D1-Cre;pDyn^+/+^) and cKO (D1-Cre;pDyn^fl/fl^) mice in these tasks suggesting that deletion of Dyn in dSPNs does not cause any gross behavioral deficits.

**Figure 5.**
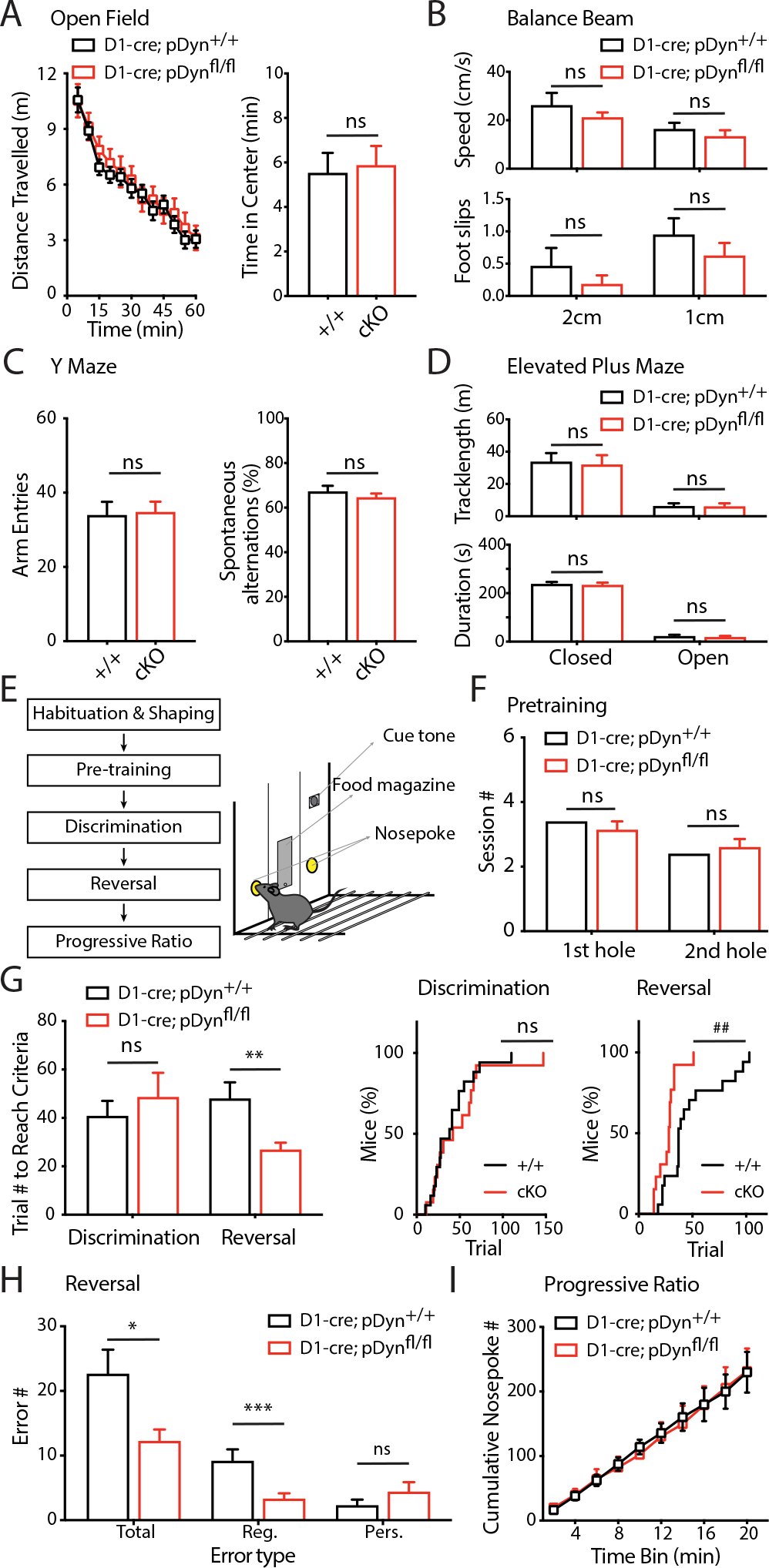
Behavioral assessment of pDyn WT and cKO mice. (**A**) Open field test. Left, distance travelled over 60 minutes (WT D1-Cre;pDyn^+/+^, n = 25; cKO D1-Cre;pDyn^fl/fl^, n = 19; genotype: F1,42 = 0.2985, p = 0.59, 2-way RM ANOVA). Right, time spent in the center (WT, 5.54 ± 0.89; cKO, 5.89 ± 0.85, p = 0.78, t test). (**B**) Balance beam test with 2 or 1 cm-diameter suspended beam. Upper, speed on 2 cm (WT, n = 7, 26.39 ± 4.93; cKO, n = 10, 21.40 ± 1.8; p = 0.37, t-test) and 1 cm beam (WT, 16.56 ± 2.42; cKO, 13.58 ± 2.34, p = 0.39, t-test). Lower, foot slips on 2 cm (WT, 0.47 ± 0.27; cKO 0.19 ± 0.13, p = 0.38, t-test) and 1 cm beam (WT, 0.96 ± 0.25; cKO, 0.63 ± 0.19, p = 0.31, t-test). (**C**) Y-maze task. Left, arm entries (WT, n = 13, 34.08 ± 3.47; cKO, n = 16, 34.88 ± 2.72, p = 0.89, t-test). Right, spontaneous alternations % (WT, 67.46 ± 2.38; cKO, 64.81 ± 1.53, p = 0.34, t-test). (**D**) Elevated plus maze test. Upper, tracklength in closed (WT, n = 25, 34.09 ± 5.03; cKO, n = 19, 32.19 ± 5.56, p = 0.8, t-test) and open arms (WT, 3.45 ± 0.76; cKO, 2.97 ± 0.74, p = 0.65, t-test). Lower, duration in closed (WT, 239.36 ± 7.19; cKO, 235.87 ± 7.59, p = 0.74, t-test) and open arms (WT, 24.46 ± 4.17; cKO, 20.23 ± 3.45, p = 0.44, t-test). (**E**) Experimental scheme and behavior apparatus of 2-choice spatial discrimination reversal learning task. (**F**) It took similar session numbers for both WT and cKO mice to learn nose-poking in pre-training stage. 1^st^ hole: WT, n = 17, 3.41 ± 0.32; cKO, n = 13, 3.15 ± 0.24. 2^nd^ hole: WT, 2.41 ± 0.15; cKO 2.62 ± 0.24, p = 0.48, t test. (**G**) Left: trial numbers the mice took to reach criteria in the discrimination (Disc) and reversal (Rev) phases. Both WT and cKO mice needed a comparable number of trials to reach the criteria during the discrimination phase (WT, 40.9 ± 6.06; cKO 48.8 ± 9.89, p = 0.51, t test), while cKO group needed significantly fewer trials to reach the criteria during the reversal phase (WT, 48.17 ± 6.56; cKO, 27 ± 2.7, p = 0.0071, t test). Right: cumulative plots showing the percentage of mice that reached criteria over training trials. The curves during the discrimination phase were not significantly different (p = 0.84, Kolmogorov-Smirnov test), while cKO reached the criteria significantly faster during the reversal phase (^##^ p = 0.0019, Kolmogorov-Smirnov test). (**H**) During the reversal phase, cKO made significantly fewer total errors (WT, 22.7 ± 3.67; cKO, 12.3 ± 1.72, p = 0.018, t-test) and specifically fewer regressive errors (WT, 9.24 ± 1.72; cKO, 3.38 ± 0.79, p = 0.00054, t-test), but comparable numbers of perseverative errors (WT, 2.35 ± 0.82; cKO, 4.46 ± 1.42, p = 0.21, t test). (**I**) In the progressive ratio task, there is no difference between WT and cKO groups in the cumulative plot of nose poke numbers (F1,23 = 0.00058, p = 0.94, 2-way RM ANOVA). And the number of rewards achieved in the session was comparable (control 15.54 ± 0.55, cKO 15.17 ± 0.89, p = 0.72, t test). Data are presented as mean ± SEM. ns: not significant, p > 0.05, * p < 0.05, ** p < 0.01, *** p < 0.001.

We then utilized instrumental reversal learning task to assess the animal’s performance during initial learning and the animal’s flexibility to adapt to a new action-reward contingency during the reversal learning phase (Izquierdo et al., 2017). We first employed a spatial reversal learning task using operant chambers with two nose-poking holes (Figure 5E; see Methods). In this task, mice needed to poke one of the two holes to receive a food reward. Mice were pre-trained to proficiently poke into both holes and to retrieve rewards from the food magazine. We found that it took both WT control (D1-Cre;pDyn^+/+^) and cKO (D1-Cre;pDyn^fl/fl^) mice similar numbers of training sessions during the nosepoke pre-training phase (Figure 5F), suggesting that pDyn cKO did not affect general action sequence learning. Next, during the spatial discrimination phase, one of the two holes was associated with reward, while the other with 5-second timeout punishment. We found that the trial numbers to reach criteria (9 correct responses out of 10 trials) during the discrimination phase was similar for both groups (Figure 5G). These results suggest pDyn cKO did not change the initial acquisition of action-reward association. In the subsequent reversal phase, the correct and wrong nosepoke holes were reversed. The mice initially poked the previously rewarded hole and were punished with a timeout, and gradually switched to the updated correct hole. Strikingly, the trial number to reach criteria during reversal was significantly reduced in the cKO (D1-Cre;pDyn^fl/fl^) group (Figure 5G). Consistently, cKO mice made significantly fewer errors (failed trials in which mice poked into the previously rewarded but currently unrewarded hole) during the reversal phase (Figure 5H). We further classified the errors into perseverative errors (failed trials before mice tried the newly rewarded hole for the first time) and regressive errors (failed trials immediately following a correct trial), which are used to measure the sensitivity to negative feedback and their flexibility in adapting to the new reward contingency, respectively (Izquierdo et al., 2017). We found that regressive errors were significantly reduced in the cKO (D1-Cre;pDyn^fl/fl^) mice, while perseverative errors were comparable (Figure 5H). These results suggested that loss of striatal dynorphin elevated the animal’s flexibility in adapting to new action-reward contingencies during the reversal phase.

Since previous studies showed that Dyn/KOR might modulate the perception of reward value (Carlezon et al., 2006; Negus, 2004; Schindler et al., 2010; Todtenkopf et al., 2004), we next examined if the enhanced performance during reversal learning was due to a change in motivation to work for the reward. We conducted a progressive ratio task with the same mice. Nosepoke rates over the entire session were indistinguishable between cKO (D1-Cre;pDyn^fl/fl^) mice and control (D1-Cre;pDyn^+/+^) mice groups (Figure 5I), suggesting that loss of striatal Dyn did not change the motivation or the perceived value of the food reward.

We further confirmed this unique behavioral phenotype with a different task - 4-choice odor discrimination reversal learning task (Johnson and Wilbrecht, 2011). A new cohort of mice was trained to dig for food reward (habituation and shaping phases) and then trained to associate a specific odor with the food reward (discrimination phase) (Figure 6A; see Methods). After mice reached the discrimination criteria (8 correct responses out of 10 trials), the food reward was paired with one of the other odors (reversal phase). Consistent with our results in the spatial reversal paradigm, we found that while the trial numbers to reach criteria in the discrimination phase were similar, the trial numbers in the reversal phase were significantly reduced in cKO (D1-Cre;pDyn^fl/fl^) mice (Figure 6B). We again quantified the errors during the reversal phase and found that cKO mice made fewer total errors, with a specific decrease in the number of regressive errors but not perseverative errors (Figure 6C), suggesting that cKO (D1-Cre;pDyn^fl/fl^) mice have a faster adaptation of the new action-reward contingency. To assess reward motivation in this task, we measured the time latency from trial starts to digging during the reversal phase and found no significant difference (Figure 6D).

**Figure 6.**
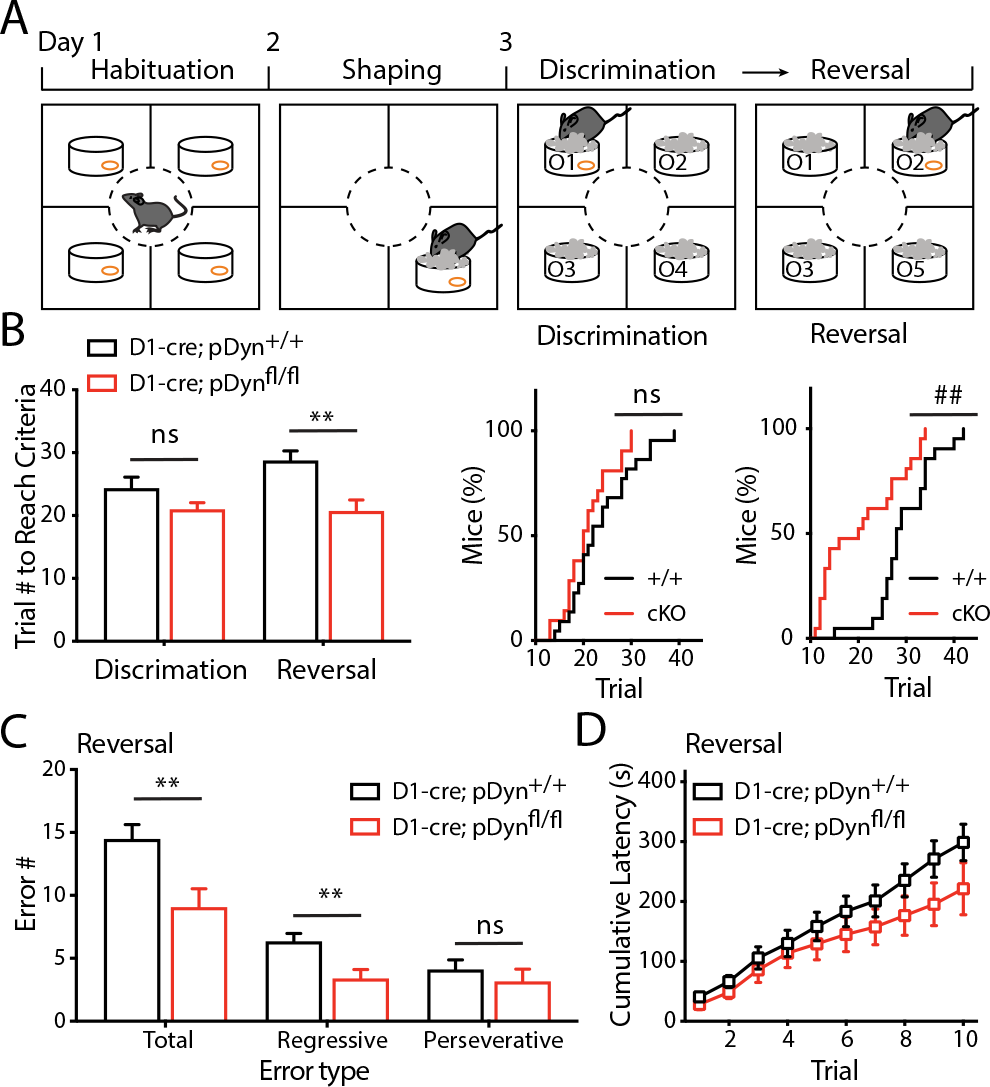
Enhanced reversal learning in pDyn cKO mice in 4-choice odor discrimination task. (**A**) Schematic of the experiment. The shavings were unscented in the shaping phase and scented with four of the five odors (O1–O5) during the discrimination and reversal phases. (**B**) Left: trial numbers the mice took to reach criteria in the discrimination (Disc) and reversal (Rev) phases. There is no significant difference in the number of trials for WT and cKO groups to reach the criteria during the discrimination (WT, n = 22, 24.32 ± 1.79; cKO, n = 21, 20.95 ± 1.09, p = 0.12, t-test), while cKO mice need significantly fewer trials to reach criteria during the reversal phase (WT, 28.73 ± 1.54; cKO, 20.67 ± 1.82, p = 0.0016, t test). Right: cumulative curves of the percentage of mice that reached criteria over training trials. The plots are not significantly different during the discrimination phase (p = 0.87, Kolmogorov-Smirnov test), and cKO group increases significantly faster than WT during the reversal phase (^##^ p = 0.0021, Kolmogorov-Smirnov test). (**C**) During the reversal phase, cKO mice made significantly fewer total errors (WT, 14.46 ± 1.17; cKO, 9.05 ± 1.48, p = 0.006, t test) and significantly fewer regressive errors (WT, 6.32 ± 0.66; cKO, 3.38 ± 0.72, p = 0.0044, t test), but similar numbers of perseverative errors (WT, 4.09 ± 0.78; cKO, 3.14 ± 1.01, p = 0.24, t test). (**D**) Cumulative plot of the time latencies from trial starts to digging over the first 10 trials in the reversal phase. 2-way RM ANOVA revealed no effect by genotype (F1,39 = 1.453, p = 0.24). Data are presented as mean ± SEM. ns: not significant, p > 0.05, ** p < 0.01.

In summary, Dyn cKO (D1-Cre;pDyn^fl/fl^) mice and control (D1-Cre;pDyn^+/+^) mice demonstrated similar performance during the initial acquisition of action-reward contingency in both discrimination reversal learning behavioral paradigms, however, the Dyn cKO mice showed increased flexibility in adapting new reward contingency during the reversal phase.

## DISCUSSION

Neuropeptides, such as Dyn, are highly expressed in unique neuron populations. They are often used as cellular markers for the identification of certain neuronal cell types. In the striatal, Dyn is highly expressed in the direct-pathway and has often been used as a cellular marker for dSPNs (Edwards et al., 2017). However, surprisingly, its endogenous functions remain unclear. In addition to the conventional axonal release, Dyn is also thought to be released through dense core vesicles located in soma and dendrites during high frequency firing (Al-Hasani et al., 2015). Because of its unique expression pattern – the ligand, Dyn, is exclusively expressed in the dSPNs, and the receptor, KOR, is also highly differentially expressed in dSPNs – Dyn/KOR has been hypothesized to dampen dSPN output, functioning similarly to an autoreceptor (Nestler, 2004). In the present study, we probed the functional role of Dyn using pathway-specific deletion of Dyn combined with pharmacology, we found an unexpected role of Dyn/KOR signaling for the regulation of striatal synaptic plasticity and behaviors. In particular, dynorphin/KOR signaling bidirectionally regulates striatal plasticity in a pathway-specific manner, together with dopamine D1R signaling, to fine tune dSPN LTP expression. We further revealed an interesting behavioral phenotype in mice lacking Dyn in dSPNs using two different discrimination reversal learning paradigms, suggesting a crucial role of striatal Dyn in cognitive flexibility.

### Dyn/KOR signaling regulates LTP in dSPNs

Striatal SPNs receive convergent glutamatergic inputs from various areas of the cerebral cortex and the thalamus, as well as the midbrain dopaminergic inputs (Gerfen and Surmeier, 2011). Previous studies have shown that Dyn can activate presynaptic KOR to inhibit glutamatergic and DA release through a Gi/o-coupled signaling cascade (Atwood et al., 2014; Di Chiara and Imperato, 1988; Ehrich et al., 2015; Mu et al., 2011; Spanagel et al., 1992). Studies from other brain regions also suggest that Dyn/KOR regulates synaptic plasticity, for example, studies in the hippocampus have shown that Dyn released from dentate granule cells blocks LTP of excitatory inputs (Drake et al., 1994; Terman et al., 1994; Wagner et al., 1993; Weisskopf et al., 1993). Similarly, in the VTA, KOR activation blocks LTP of inhibitory inputs onto dopaminergic cells (Graziane et al., 2013; Polter et al., 2014).

Here, we show Dyn/KOR signaling bi-directionally regulate LTP expression in dSPNs using both genetic and pharmacological tools. A previous study reported that acutely evoked release of Dyn suppress LTP in dorsal striatum indirectly via reducing striatal dopamine levels (Hawes et al., 2017). It is certainly true that acute KOR activation alters DA levels in the *ex vivo* slice. However, we demonstrated that D1-pDyn cKO did not affect dopaminergic basal transmission, therefore, an altered DA release is unlikely responsible for bi-directional regulation of dSPN LTP. KORs are expressed in both pre- and postsynaptic sites (Svingos et al., 2001). In addition to its prominent presynaptic actions on glutamatergic and dopaminergic inputs to SPNs (Nestler, 2004), postsynaptic KOR activation can also lead to AMPARs endocytosis via protein kinase A (PKA)/calcineurin-dependent mechanisms in interneurons (Coleman et al., 2021). Because LTP expression in dSPN requires postsynaptic activation of D1R, which subsequently upregulates the adenylate cyclase (AC)/protein kinase A (PKA) cascade pathway (Surmeier et al., 2014), and KOR can downregulate AC/PKA signaling through Gi/o-coupled mechanism (Bruchas and Chavkin, 2010; Gentleman et al., 1983), it is likely that Dyn/KOR signaling directly interacts with D1R signaling to bi-directionally regulate LTP expression in dSPNs through a postsynaptic mechanism.

### Interaction between Dyn/KOR and dopaminergic D1R signaling

Previous studies established a model in which different GPCR cascades modulate synaptic plasticity in SPNs (Surmeier et al., 2014). In particular, D1R activation is required to enable Hebbian LTP induction, and deactivating D1R blocks LTP and shift the plasticity polarity to LTD (Shen et al., 2008). Our current study adds an additional player - Dyn/KOR signaling - to the model. Gs-coupled D1R signaling and Gi/o-coupled KOR signaling pathways converge and bi-directionally regulate AC/PKA levels in opposing directions. However, the relative strength is subtly different. We show that when D1R is fully activated LTP expression is enhanced regardless of whether Dyn/KOR signaling is activated or blocked, supporting the model that D1R activation is permissive for LTP expression, and high levels of D1R activation can saturate LTP expression machinery, thus occluding the effect of pDyn cKO or KOR blockade. Conversely, Dyn/KOR signaling may exert a more prominent effect under lower levels of D1R activation (i.e., the basal slice condition or with D1R blockade). Under the basal slice conditions without pharmacological D1R manipulations, blocking KOR and pDyn cKO both enhanced LTP expression, and KOR activation abolished LTP expression. Remarkably, blocking KOR or deleting Dyn both were able to rescue LTP expression even when D1R is fully blocked. In psychomotor disorders, such as the Parkinson’s disease, in which the dopamine depletion leads to a lack of LTP in dSPNs (Pisani et al., 2005; Surmeier et al., 2014) and thus a disrupted balance of the striatal network. In addition to dopamine replacement therapy to restore dopamine receptor signaling, KOR antagonism may hold potential to rescue striatal plasticity and alleviate striatal circuit dysfunctions.

### Striatal Dyn impedes cognitive flexibility during learning

Dyn in the central nervous system has long been recognized as a major mediator of dysphoria, anxiety, and depression during stress and drug withdrawal (Crowley et al., 2016; Land et al., 2008; Van’t Veer and Carlezon, 2013). Constitutive knockout of dynorphin reduces anxiety and depression levels (Schwarzer, 2009), and evoked dynorphin release *in vivo* by optogenetic activation of dSPNs leads to aversion (Al-Hasani et al., 2015; Soares-Cunha et al., 2020). The striatal direct-pathway pDyn cKO mice showed normal anxiety levels, suggesting the reduced anxiety phenotype may be mediated by other brain regions (Crowley et al., 2016).

By contrast, studies focusing on learning and memory have generated conflicting results (Carey et al., 2009; Hiramatsu and Hoshino, 2004; Kuzmin et al., 2006; Ukai et al., 1997). Such discrepancy may result from the widespread expression of Dyn and KOR, and the functional complexity of Dyn/KOR signaling across various brain regions. Therefore, it is crucial to study Dyn function using cell-type-specific approaches. Here, we unexpectedly identified enhanced cognitive flexibility phenotype of striatal pDyn cKO mice during reversal learning. Our finding is consistent with previous studies that reported the neurotoxic lesions of the dorsal striatum impaired various reversal learning tasks (Braun and Hauber, 2011; Castane et al., 2010). In addition, dSPNs were shown to be particularly relevant for learning flexibility (Darvas and Palmiter, 2015). Here, we provide mechanistic insights linking striatal Dyn/KOR signaling, synaptic plasticity, and flexible reversal-learning behavior.

We speculate that the interaction between Dyn/KOR and D1R signaling may contribute to the behavioral phenomenon we revealed here that D1-pDyn cKO specifically showed enhanced learning specifically in the reversal phase while not affecting the initial phase. Phasic dopamine transients in response to reward-prediction error act as the teaching signaling during the reward learning (Watabe-Uchida et al., 2017), and D1R activations are highly dependent on the phasic dopamine release due to its relatively low ligand-binding affinity (Richfield et al., 1989). Given that the suppression of LTP by Dyn/KOR is masked by high D1R activation and revealed under low D1R activation (Figure 4), it is possible that the dopamine levels are different during the initial learning phase and the reversal phase, and the relatively lower dopamine levels during the reversal phase allowed the enhancement of LTP caused by Dyn deletion and enhanced cognitive flexibility in Dyn cKO mice.

Our results also imply that striatal Dyn release may impede cognitive flexibility. While this trait seems disadvantageous for animals’ survival in a changing environment, ethological studies showed a potential trade-off between the ability to rapidly acquire new memories and the ability to retain old memories (Lea et al., 2020; Tello-Ramos et al., 2019). For instance, many food-caching species that displayed long-lasting spatial memory performed significantly worse than their closely related non-caching species during reversal learning. Therefore, striatal dynorphin may contribute to long-term memory retention.

## ACKNOWLEDGEMENTS

We thank Dr. Eddy Albarran, Dr. Richard Roth for advice on the manuscript, Drs. Yu Liu and Xiaobai Ren for technical assistance, and the members of Ding lab for valuable discussions. This work was funded by NINDS/NIH NS091144 (to J.B.D.), the GG gift fund (to J.B.D.), the Stanford Bio-X Bowes Graduate Student Fellowship (to R.Y.), and Stanford Neuroscience Institute Interdisciplinary Scholar Awards (to R.R.L.)

## AUTHOR CONTRIBUTIONS

R.Y., R.R.L., and J.B.D. conceived this project. R.Y. performed electrophysiology recording and operant chamber behavior experiments. R.Y. and R.R.L. analyzed the data. R.R.L. performed immunostaining and behavioral experiments. F.J.H. performed the spine imaging. D.K. and J.B.D. designed pDyn targeting constructs. R.Y., D.W.B. and J.B.D. wrote the manuscript with input from all authors.

## DECLARATION OF INTERESTS

The authors declare no competing interests.

## STAR٭METHODS

### KEY RESOURCE TABLE

| REAGENT or RESOURCE | SOURCE | IDENTIFIER |
| --- | --- | --- |
| **Antibodies** |  |  |
| Rabbit anti-dyn antisera | Peninsula Labs | Cat# T4268 |
| Goat anti-rabbit IgG, Alexa Fluor 488 conjugate | Invitrogen | Cat# A11008 |
| **Chemicals, Peptides, and Recombinant Proteins** |  |  |
| VU10010 | Tocris | Cat# 3141 |
| (-)-U50488 | Tocris | Cat# 0495 |
| Colchicine | Tocris | Cat# 1364 |
| Nor-binaltorphimine (NorBNI) | Tocris | Cat# 0374 |
| SCH23390 | Tocris | Cat# 0925 |
| SKF81297 | Tocris | Cat# 1447 |
| Gramicidin A | Sigma-Aldrich | CAS# 11029-61-1 |
| Alexa Fluor 594 | Invitrogen | Cat# A10438 |
| Tetrodotoxin citrate | A.G. Scientific | Cat# T-2785 |
| Picrotoxin | Tocris | Cat# 1128 |
| **Experimental Models: Organisms/Strains** |  |  |
| Mouse: pDyn<sup>fl/fl</sup> | Generated in the lab | N/A |
| Mouse: C57BL/6J | The Jackson Laboratory | Cat # 000664 |
| Mouse: Drd1-cre | MMRRC | Cat # 017264-UCD |
| Mouse: Ai32 | The Jackson Laboratory | Cat # 012569 |
| **Software and Algorithms** |  |  |
| ImageJ | NIH | RRID: SCR_003070 |
| MATLAB | MathWorks | RRID: SCR_001622 |
| GraphPad Prism 6 | GraphPad Software | RRID: SCR_002798 |

### CONTACT FOR REAGENT AND RESOURCE SHARING

Further information and requests for resources should be directed to and will be fulfilled by the Lead Contact, Jun Ding.

### EXPERIMENTAL MODEL AND SUBJECT DETAILS

#### Mice

Mice were housed under a 12 h light/dark cycle and water and food were available ad libitum. Animal experiments were conducted following protocols approved by Administrative Panel on Laboratory Animal Care at Stanford University. pDyn^fl/fl^ mice were generated by Stanford Transgenic, Knockout and Tumor model Center (TKTC). Double or triple transgenic mice were generated by crossing pDyn^fl/fl^ with D1-cre mice (MMRRC stock # 017264-UCD) (Gong et al., 2007), and Rosa-CAG-LSL-ChR2(H134R)-eYFP mice (Ai32, purchased from JAX #012569). For electrophysiological studies, D1-Cre;pDyn^+/+^;Ai32 were used as wildtype (WT) control mice to compare with the dSPN Dyn cKO (D1-Cre;pDyn^fl/fl^;Ai32) mice. The expression of ChR2 allowed us to optically identify dSPN during the recording. Neurons with SPN intrinsic properties and failed to respond to blue light stimulation (450nm, 5-10 mW) were identified as iSPNs. For electrophysiology, both male and female (4-12 weeks) mice were used. For behavioral tests, only D1-Cre;pDyn^+/+^ (WT control) and D1-Cre;pDyn^fl/fl^(cKO) without Ai32 allele were used. Both male and female adult (8-16 weeks) mice were used for behavioral tests. We used the same cohort of mice for the open field test, Y maze, three-chamber social interaction test, and elevated plus maze test; one cohort for 4-choice behavior, and another cohort for spatial reversal learning. During 4-choice behavior and spatial reversal-learning task, mice started to be food-restricted 3 days before the tasks and monitored daily to maintain at 90% body weight.

### METHOD DETAILS

#### Generation of pDyn cKO Mice

The murine prodynorphin gene (pDyn) contains 4 exons. A BAC clone containing exon 4 genomic sequence was used to generate the loxP-flanked pDyn targeting construct. A PCR amplicon containing the loxP-NEO/Kan-loxP cassette from the pSV-Cre plasmid flanked by 70 bp of homologous sequence at both ends, which matched sites in the third intron of pDyn, was generated using the following primers: DYN-LOXP-F1, 5’ TGTAAGCGCAGGGCAATCGGAAGATCTTGCTTCTACTTTTGGCCTTGCCACCCGTGTCTCTGCGCCTTTGGCAGCCCAATTCCGATCATATTCAATAACC 3’ and DYN-LOXP-R1, 5’ CTGAAAGGATGCAAGAGTTTTGTCATGTGAGAAATGAGAAATTAGACTTACAGAGCCTAGGCAGCTATCCCGCTCTAGAACTAGTGGATCCCCTCGAGGGACC 3’. The PCR amplicon was transformed into electrocompetent EL350 cells that had been previously transformed with the aforementioned pDyn BAC, followed by arabinose-induced expression of cre recombinase to remove LoxP-NEO/Kan-LoxP selective cassette (Lee et al., 2001). The modified pDyn BAC containing a loxP site in the third intron was then transformed into EL250 cells. A second PCR amplicon containing the FRT-NEO/Kan-FRT-loxP flanked by 70 bp of homologous sequence at both ends, which matched sites in the 3’-end downstream genomic region of pDyn, was generated using the following primers: DYN-LOXP-F2, 5’ ACATGTATAGCTGTATTGGAGGATTGAACCTAGTTCCTCATGCATGATAGGCAAACACTCTACCACCAAGC-CGACGGTATCGATAAGCTTGATATCGAATTCC 3’ and DYN-LOXP-R2, 5’ ACACTTGCTTTCTTGCTTTCTGTTATTCCACACTGTATTTTCTAATATGACTACAGGGGCTGGAGCTATG-GCTCTAGAACTAGTGGATCCACCTAATAACTTC 3’. The PCR amplicon was then transformed into EL250 cells containing the modified pDyn BAC. A targeting construct was derived from this modified BAC spanning the region 5 kb upstream of the loxP site inserted into intron 3 and 2 kb downstream of the loxP site inserted into 3’-end genomic region. The resulting targeting construct was then electroporated into mouse embryonic stem (ES) cells (W4/129S6, Taconic, NY). Correctly targeted ES clones were identified and injected into blastocysts resulting in several 100% male chimeras. Chimeras were bred to mice bearing a flp-recombinase transgene (ROSA-Flp, JAX #003946) to remove the neomycin selection marker, generating mice with one allele of pDyn flanked by loxP sites at exon 4 (pDyn^fl/+^). The resulted pDyn^fl/+^ offspring were then backcrossed with C57BL/6 mice for seven generations. The following primers were used for genotyping PCR setups: pDyn-F: 5’ CATGCCTGATGAAGGTCGGTAG 3’; pDyn-R: 5’ CATCTTCCAAGTCATCCTTGCC3’; and pDyn-D: 5’ GCAGTACCTCTCTCTGTCCTC3’, with amplified PCR products of 315 bp for wildtype alleles, 420 bp for loxP-flanked alleles, and 552 bp for cre-deleted alleles. We then crossed pDyn^fl/+^ with D1-cre mice (MMRRC stock # 017264-UCD) (Gong et al., 2007) to obtain D1-cre;pDyn^fl/+^ mice. For experiments, dynorphin conditional knockout mice (D1-cre; pDyn^fl/fl^) and littermate controls (D1-cre; pDyn^+/+^) were obtained from breeding pairs of D1-cre; pDyn^fl/+^ × pDyn^fl/+^.

#### Immunohistochemistry

Immunohistochemistry was performed as described in (Wu et al., 2015). To stain for dynorphin, mice were intracerebroventricularly injected with the axonal transport blocker colchicine (75 µg/kg body weight) 48 hours before procedures to reveal dynorphin immunoreactivity within neuronal cell bodies (Li and van den Pol, 2006). Mice were anesthetized with isoflurane and intracardially perfused with ice-cold 4% paraformaldehyde in phosphate buffer (PB). Brains were dissected, post-fixed 24 h at 4 °C and cryoprotected with a solution of 30% sucrose in 0.1 M PB at 4 °C for at least 24 h, cut into 30-µm sections and processed for immunostaining. Sections were washed three times in PBS and blocked in PBS containing 0.5% Triton X-100 (G-Biosciences) and 5% normal goat serum (Cell Signaling). Sections were then incubated overnight at 4 °C with rabbit anti-dyn antisera (1:1000, Peninsula Labs). Following incubation, sections were washed three times in PBS and then incubated for 2 h at room temperature in Alexa488 goat anti-rabbit IgG (1:1000, Invitrogen). Sections were then washed three times in PBS. This was followed by a 1-h incubation with fluorescent DAPI stain to allow visualization of cell bodies (1:400, Neurotrace, Invitrogen). Sections were then washed three times in PBS, followed by three 10-min rinses in PB and mounted on glass slides with Hard set Vectashield (Vector Labs) for microscopy. All the sections were imaged using confocal microscopy (Leica TCS SPE confocal microscope) with consistent settings. Cells were quantified in ImageJ.

#### Electrophysiology

##### 1. Slice preparation

Brain slices (300 μm) were obtained from 2-4-month-old mice using standard techniques (Wu et al., 2015). Briefly, animals were anesthetized with isoflurane and decapitated. The brain was exposed and chilled with ice-cold artificial CSF (ACSF) containing 125 mM NaCl, 2.5 mM KCl, 2 mM CaCl2, 1.25 mM NaH2PO4, 1 mM MgCl2, 25 mM NaHCO3, and 15 mM D-glucose. ACSF was saturated with 95% O2 and 5% CO2. Osmolarity was adjusted to 300-305 mOsm. Oblique horizontal brain slices containing the dorsomedial striatum were prepared with a vibrating microtome (Leica VT1200 S, Germany) and left to recover in ACSF for 30 min at 34°C and then at room temperature for an additional 30 min before recording. For pharmacological treatment with NorBNI (1 µM), U50488 (1 µM), SCH23390 (3 µM), SKF81297 (3 µM), and VU10010 (5µM), the ACSF for recovering also contains the corresponding drugs. After the recovery period, slices were moved to a submerged recording chamber perfused with ACSF at a rate of 2-3 ml/min at 30-31°C, and brain slices were recorded within 5 hours after recovery.

##### 2. Fast-scan cyclic voltammetry

Brain slices from D1-cre;pDyn^+/+^ and D1-cre;pDyn^fl/fl^ mice were used for fast-scan cyclic voltammetry recordings to measure the extracellular dopamine release. Recordings were performed using carbon-fiber microelectrodes (7 μm diameter carbon fiber extending 50-100 μm beyond the tapered glass seal). Cyclic voltammograms were measured with a triangular potential waveform (-0.4 to +1.3 V versus Ag/AgCl reference electrode, 400 V/s scan rate, 8.5 ms waveform width) applied at 100 ms intervals. The carbon fiber microelectrode was held at -0.4 V between scans. Cyclic voltammograms were background-subtracted by averaging 10 background scans. Dopaminergic axon terminals were stimulated locally (100-200 μm from carbon fiber) with a concentric bipolar electrode (FHC) with a single stimulation (100 μA intensity, 0.2 ms length) or a train of three stimulations in 50 Hz. Recordings were obtained using TarHeel CV system. Changes in dopamine concentration by electrode stimulation were quantified by plotting the peak oxidation current of the voltammogram over time. Carbon-fiber microelectrode was calibrated at the end of each day of experiments to convert oxidation current to dopamine concentration using 10 μM dopamine in ACSF.

##### 3. miniature EPSC recording

Brain slices from D1-cre;pDyn^+/+^;Ai32 and D1-cre;pDyn^fl/fl^;Ai32 mice were used to record miniature EPSCs (mEPSCs). SPNs were visualized under infrared illumination using an Olympus BX51WI microscope equipped with DIC, a water-immersion objective (40× NA 0.8), and a CMOS camera (Hamamatsu Photonics). Whole-cell voltage-clamp recording was performed with borosilicate glass microelectrodes (3-5 MΩ) filled with a Cs^+^-based low Cl^-^ internal solution (126 mM CsMeSO3, 8 mM NaCl, 10 mM HEPES, 2.9 mM QX-314, 8 mM Na2-Phosphocreatine, 0.3 mM GTP-Na, 4 mM ATP-Mg, 0.1 mM CaCl2, 1 mM EGTA; pH 7.2-7.3; osmolarity 285-290 mOsm). The access resistance was < 25MΩ (no compensation), and the data were discarded if the access resistance changed more than 25% during recording. The presence of ChR2 current evoked by a 450 nm laser pulse (0.5 ms, OptoEngine, USA) was used to distinguish dSPNs and iSPNs. For recording of miniature EPSC (mEPSC), TTX (1 µM) was included, and neurons were held at a membrane potential of the Cl^-^ reversal potential (-70 mV, liquid junction potential not corrected). Recordings were obtained with a Multiclamp 700B amplifier (Molecular Devices) using the WinWCP software (University of Strathclyde, UK). Signals were filtered at 2 kHz, digitized at 10 kHz (NI PCIe-6259, National Instruments), and analyzed offline using Clampfit 10.0 (Molecular Devices) and Mini Analysis Program (Synaptosoft).

##### 4. Perforated patch and LTP induction

Brain slices from D1-cre;pDyn^+/+^;Ai32 and D1-cre;pDyn^fl/fl^;Ai32 were used to record LTP. The perforated patch and LTP induction protocol was adapted from a previous study (Shen et al., 2008). Electrical access was achieved through the perforated-patch method using Gramicidin A (Sigma). Perforated patch was performed with a borosilicate glass microelectrode (3-3.5 MΩ), front-filled with 1 µl K^+^-based internal solution (135 mM KMeSO3, 8.1 mM KCl, 10 mM HEPES, 8 mM Na2-Phosphocreatine, 0.3 mM GTP-Na, 4 mM ATP-Mg, 0.1 mM CaCl2, 1 mM EGTA; pH 7.2-7.3; osmolarity 285-290 mOsm), and back-filled with 10 µl Gramicidin A-containing internal solution. The Gramicidin A-containing internal solution was made fresh before use: a stock solution of Gramicidin A (20 mg/mL) in dimethyl sulfoxide (DMSO) was prepared and diluted in the K^+^-based internal solution yielding a final concentration of 200 µg/mL. The fluorescent dye Alexa594 (10 µM) was also added in the internal solution for visualizing the integrity of the perforated patch configuration throughout the recording. After the microelectrode formed a giga seal with the cell membrane, access resistance was continuously monitored during perforation by applying a -5 mV pulse from a holding potential of -70 mV, under the voltage-clamp mode. A stable perforated patch normally formed within 30-60 minutes, and the access resistance stays around 30-50 MΩ without further decreasing. Subsequently, the presence of ChR2 current evoked by a 450 nm laser pulse (0.5 ms, OptoEngine, USA) was used to distinguish dSPNs from iSPNs. Then the recording was switched to current-clamp mode, and serial resistance was compensated with amplifier bridge balance. The data were excluded if the input resistance (Rin) changed more than 25% over the course of the experiment. Presynaptic inputs, recorded as EPSPs, were evoked by focal extracellular stimulation with a small theta glass electrode positioned 50-100 µm from the recorded cell body. Stimulation intensity (0.2 ms, 5-30 µA) was adjusted to evoke stable EPSPs with an amplitude of around 2-5 mV. EPSPs were evoked every 20 s for 5 minutes as the baseline. Then LTP was induced using the spike-timing-dependent plasticity (STDP) protocol. The protocol consisted of 15 trains of five bursts repeated at 0.1 Hz, with each burst composed of three postsynaptic spikings preceded (+5 ms) with three presynaptic inputs (EPSPs) at 50 Hz (Figure 5A). The postsynaptic spikes were evoked by direct somatic current injection (5 ms, 1-1.5 nA). During the induction, the postsynaptic cell membrane potential was depolarized to -70 mV. After the STDP induction, EPSPs were recorded for another 30 minutes to monitor the change of amplitudes. GABAA were blocked by the bath application of 100 µM picrotoxin throughout the recording. For recordings with NorBNI (1 µM), (-)-U50488 (1 µM), SCH23390 (3 µM), or SKF81297 (3 µM), or VU10010 (5µM), slices were incubated with these drugs during slice recovery (one hour) as well as throughout the recording, which is to reduce the acute effect of a receptor agonist or antagonist treatment.

#### Behavioral assays

##### 1. Open field test

Mice were individually placed in the center of illuminated white plastic boxes (40 × 40 × 40 cm) and allowed to roam freely for 60 min. Locomotion was video recorded and analyzed using Viewer III tracking system (Biobserve). Total distance traveled and time spent in the 20 × 20 cm center of the square were quantified. The total distance was used to evaluate locomotor activity and the time spent in center area was used to estimate the anxiety level in an open environment. The total distance was quantified over each 5-min bin and analyzed with repeated measurement two-way ANOVA. The time spent in the center were analyzed with Student’s t-test.

##### 2. Balance beam test

The animal’s ability to navigate across a beam was tested using two wooden beams (cylindrical beam 95 cm in length, 1 cm and 2 cm in diameter, respectively). The beams were fixed with ring stands on both ends 50 cm above the ground with one end leading into a small hide box. During the trials, the 2 cm beam was first used. Mice were placed at the start line near one end of the beam, facing the hide box, which was 80 cm away to the other end. Mice were left to walk across the beam to the box. After reaching the hide box, mice were let to stay there for 1 min and placed at the start line again for a total of three trials. Repeat the trials with 1 cm beam. The speed, number of foot slips for each mouse were scored and averaged across 3 trials for each beam. Data were analyzed with Student’s t-test.

##### 3. Spontaneous Y maze

The Y-maze contained three gray arms (each arm: 40 × 10 × 17 cm) placed at 120° angles. Mice were placed in the distal end of one arm and allowed to freely explore the maze for 10 min under video monitoring. One entry was counted when the whole body including tail of a mouse entered an arm. An alternation was defined when the mouse visited all three different arms in three consecutive entries. The alternation percentage was calculated as [number of alternations/total number of entries] × 100. Y maze is based on the innate tendency of rodents to explore novel environment; thus, a mouse would try to make more spontaneous alternations in the Y maze, which depends on its spatial working memory. The alternation percentage was used to evaluate the spatial working memory of mice. Data were analyzed with Student’s t-test.

##### 4. Elevated plus maze

The maze was elevated 40 cm above the floor and consisted of two open arms without walls and two closed arms with 10-cm tall walls. Each arm was 30 × 8 cm, connected by an 8 × 8 cm center area. Mice were habituated in the testing room 1 hour before testing. Mice were placed in the center zone of the EPM facing one of the open arms and were allowed to explore freely for 10 minutes. Locomotion was video recorded and analyzed using Viewer III tracking system (Biobserve). Total distance traveled, time and distance spent in the closed and open arms were quantified. Data were analyzed with Student’s t-test.

##### 5. Three-chamber sociability test

A transparent three-chamber apparatus (60 × 30 × 30 cm per chamber) was used for sociability tests. One day before testing, the mice were first habituated in the entire apparatus for 20 minutes. During testing, a stranger C57BL/6J mouse confined in an upside-down wired-mesh cup was placed in one chamber (social chamber), and an identical but empty cup was placed in the right chamber (non-social chamber). The mice were placed in the center chamber and left to freely explore the three chambers for 30 min. The activity was recorded with Viewer III tracking system, and time spent in each chamber was quantified. Data was analyzed with Student’s t-test.

##### 6. Spatial reversal learning task

The training task was adapted and modified from previous studies (Castane et al., 2010; Ineichen et al., 2012). The training was conducted daily Monday-Sunday. All mice underwent only one training session per day throughout the whole task. The mice started to be food restricted 3 days before the training and monitored to maintain around 90% body weight.

###### 6.1. Operant apparatus

Each operant chamber was measured 21.6 × 17.8 × 12.7 cm (model # ENV-307W; Med Associates) and housed within a sound and light attenuating cubicle (ENV-022MD; Med Associates). The back and ceiling and a forward-opening door were transparent plexiglass, and the floor was stainless steel grid. A pellet dispenser delivering a reward pellet (20 mg, #F06649; BioServ) singly into a food magazine was located at one side wall, and the nose-poke ports were situated left and right of the food magazine. Each nose-poke port had a white lamp in the recess. The nose pokes and pellet retrievals were detected via infra-red beam. A house light and tone generator were located on the other side wall. A house light was provided during the training and task sessions. All elements were removed and cleaned with 70% ethanol after each session. Four such operant chambers were run in parallel, with two chambers for males and females, respectively. Each mouse was assigned to a specific chamber. The four chambers were controlled from one PC using Med-PC V interface (Med Associates). A customized program scripted in Trans V was run to automatically control and record all experimental events and outputs. Only one session was conducted for each mouse per day.

###### 6.2. Habituation and shaping

Mice were taken to the chambers for habituation for 3 days. The house light was lighted, and food magazine was loaded with pellets, and mice were placed in the operant chambers individually to explore freely for 10 minutes. This was to habituate the mice to the chamber and to retrieving pellets out of the food magazine.

During shaping, the session started with turning on the house light, and a pellet was dispensed along with a 2-s tone (CS, conditioned stimulus) every 45s. Total 39 pellets were dispensed in the 30-minute session. This step was to condition the mice to associate reward pellets with the tone, which facilitated later trainings. The criteria for each mouse to proceed to next stage was retrieving at least 30 pellets for two consecutive sessions.

###### 6.3. Pre-training/nose-poke learning

For all subsequent training stages, some common procedures were kept consistent as below:

1. During each trial, nose-poke port lights were illuminated to signal the beginning of a trial.
2. Immediately after each correct nose poke, the 2-s CS tone was presented, the nose-poke port lights were shut off, and a reward pellet was delivered. Before the mouse retrieved the pellet from the magazine, further nose pokes had no consequence. After the mouse retrieved the pellet, an inter-trial-interval (ITI) of 2.5s was initiated, during which the nose-poke port lights remained off, and nose pokes had no consequence. The onset of the next trial was signaled by switching on nose-poke port.
3. After mice poking into the wrong port, a timeout punishment of ITI of 5.0 s was initiated, during which the nose-poke port lights were switched off and further nosepokes had no consequences. After 5 s, the nose-poke port lights were switched on automatically to signal the new trial.

The pre-training was to get mice familiar with the operant task of nose poking to get reward. Training took place on each port separately with FR1 schedule. Only one of nose poke ports was presented and illuminated (so there was no wrong poke). Crushed pellets were initially placed in the port to encourage the mice to poke. One poke would trigger the CS tone and pellet reward delivery. The criterion was to reach 30 pellets in two consecutive sessions. Each session lasts up to 30 minutes or when the mice reached the criteria earlier. If the criterion was not achieved, the same training was repeated the next day. After mice reached criteria for one port, the same training took place in the other port. The order of left and right ports was counterbalanced across subjects.

###### 6.4. Acquisition of spatial discrimination

After the mice learned the operant task of nose poking into one port, they were now presented with two illuminated ports, however, with one port as correct and the other as wrong. The assignment of left or right port as active was counterbalanced across subjects. As described above, in each trial, the correct poke led to reward delivery, while the wrong poke led to 5 s timeout punishment. The criteria to complete the acquisition phase was to have 9 correct responses in 10 consecutive trials in a session. Each session lasted up to 30 minutes or when the mice reached the criteria earlier. The main variables analyzed were the total number of trials and errors (failed trials) to reach criteria in the Acquisition stage. Errors made were refined by looking specifically at “perseverative” and “regressive” errors. Perseverative errors were defined as all the errors before the first correct trial. Regressive errors were those errors right after a correct trial. Some mice took more than one session to reach the criteria, in this case, the variables described above from all sessions were combined. To proceed to the next stage, mice needed to complete another “retention” session, to reinforce the acquisition of spatial discrimination.

###### 6.5. Reversal learning

The reversal learning task examined the differences in behavioral flexibility between control and pDyn cKO mice. During the task, correct and wrong nose poke ports were switched, while the other procedures remained the same with the discrimination phase. The learning criteria was the same as discrimination phase, 9 correct responses out of 10 consecutive trials. And the total number of trials, errors, as well as perseverative and regressive errors were analyzed.

###### 6.6. Progressive ratio schedule task

After the mice completed reversal learning task, they underwent progressive ratio schedule task once to assess the differences in incentive-motivation between control and pDyn cKO mice.

Only one nose-poke port was deployed and illuminated. The reinforcement was a 2-s CS tone and a pellet, the same as above, but the nose-poke port light remained on. The number of nose pokes required to get a reinforcement reward was based on exponential progression derived from the formula (5 × e^0.2n^) −5, where n was the position in the ratio sequence (Richardson and Roberts, 1996). The session lasted up to 60 minutes or after 5 min elapsed without any responses. The nosepoke rate over the whole session and reward number achieved were analyzed.

##### 7. The 4-choice odor discrimination and reversal task

The task followed the protocols from a previous study (Johnson and Wilbrecht, 2011). The task was conducted daily for three days. Mice were food deprived over 3 days before the task to 90% of their original body weight and fed to maintain 90% body weight during training.

###### 7.1. Apparatus

The 4-choice arena was a square clear acrylic box (12ʺ × 12ʺ × 9ʺ). The arena was partially divided into 4 quadrants by four 3” wide inner walls (Figure 4.a). A white ceramic bowl (2.88” in diameter, 1.75” deep) was placed in each quadrant to hold scented wood shavings to present the odors, and a Honey Nut Cheerio piece (General Mills, Minneapolis, MN) buried in the shavings as reward. A removable 6” diameter cylinder was used to confine the mice in the center between trials.

###### 7.2. Habituation

The first day of training was a habituation phase that allowed mice to become familiar with the arena and bowls containing Cheerio. A bowl containing 10 mg cheerio (no shavings) was placed in each quadrant. A mouse was placed in the cylinder. The cylinder was then lifted to let the mouse explore freely and eat from the bowls for 10 minutes. After 10 minutes, the mouse was returned to the start cylinder and bowls were re-baited. This procedure was repeated twice for a total habituation time of 30 min. The maze was wiped with 70% ethanol between animals.

###### 7.3. Shaping

The second day of training was a shaping phase to train mice to dig wood shavings for cheerio pieces. Each mouse underwent 12 training trials. In each trial, one of the four quadrants was set up with a bowl containing 10 mg cheerio buried with wood shavings. The quadrant holding the bowl was rotated over trials (SE to NW to SW to NE). And the wood shaving amounts were increased over trials: the first two with no shavings, the next two with a thin layer, then two with a quarter full, two with half full, and four trials with the bowl full of shavings burying the cheerio. Trials were not timed.

###### 7.4. Acquisition of odor discrimination

The third day consisted of an odor discrimination phase and a reversal phase. In the odor discrimination phase, mice had to discriminated four odors and learn which odor was associated with cheerio reward. Wood shavings scented with odors (O1-O4: anise, clove, litsea and thyme, respectively) were placed in separate bowls. Only O1 was rewarded with 10 mg cheerio buried. To avoid spatial association, the four odors were randomly placed in different quadrants and an odor was never in the same location two trials in a row. The mouse was first confine in the center cylinder, and the trial began when the cylinder was lifted. The mouse could freely explore the quadrants until it began digging in one of the bowls, which was immediately followed by the lowering of the cylinder to prevent multiple digging choices. The digging was defined as consistent moving shavings with both front paws, but not as sniffing or chewing shavings. When the mouse began digging in an incorrect odor, the trial was terminated, and the mouse was returned to the center cylinder; if the mouse began digging in the correct odor, it was allowed to retrieve the reward before returned to the center cylinder. If the mouse didn’t make a digging choice within 3 minutes, the trial was terminated and recorded as an omission. Analysis of the latency to dig did not include omissions. Between each trial, the odors were rearranged and re-baited if necessary. The criterion for this phase was 8 correct digging responses out of 10 consecutive trials, including the omissions.

###### 7.5. Reversal learning

Once criterion was met for discrimination, the reversal phase was initiated immediately within the same session. In this phase, O2 (clove) became the rewarded odor and O4 (thyme) was switched out for a novel odor O5 (eucalyptus). The criterion to complete this phase was the same as discrimination: 8 correct digging responses out of 10 consecutive trials. To quantify the reversal learning phenotype, their number of trials and errors before reaching criterion was analyzed. Errors were refined as follows: (1) Reversal – mouse dug in previously rewarded odor (anise); (2) Perseverative – number of reversal errors made before one correct trial; (3) Regressive – reversal errors made after one correct trial. To analyze their motivational aspects during the reversal learning, the time latency from trial starts to digging was analyzed.

## QUANTIFICATION AND STATISTICAL ANALYSIS

Statistical analyses were performed using GraphPad Prism (GraphPad Software) and MATLAB (Mathworks). Most behavioral performances between WT control and cKO groups were compared with Student’s unpaired t-test and repeated measurement 2-way ANOVA. The cumulative plot of the percentage of mice reaching the criteria were compared with the Kolmogorov-Smirnov test. Averaged mEPSC amplitudes and frequencies, spine densities, and evoked [DA]_o_ were analyzed with the Mann-Whitney Rank Sum test, and cumulative distribution of mEPSC amplitudes and inter-event intervals were analyzed with the Kolmogorov-Smirnov test. The EPSP amplitudes at the baseline and the end of recordings in LTP experiments were compared with Paired Wilcoxon signed-rank test to determine if LTP or LTD were generated, and the normalized End EPSP amplitudes were compared between conditions using the Mann-Whitney Rank Sum test. All data were represented as mean ± SEM. Within the figures: * p < 0.05, ** p < 0.01, *** p < 0.001 and ns = not significant, p > 0.05.

---

**Figure S1.**
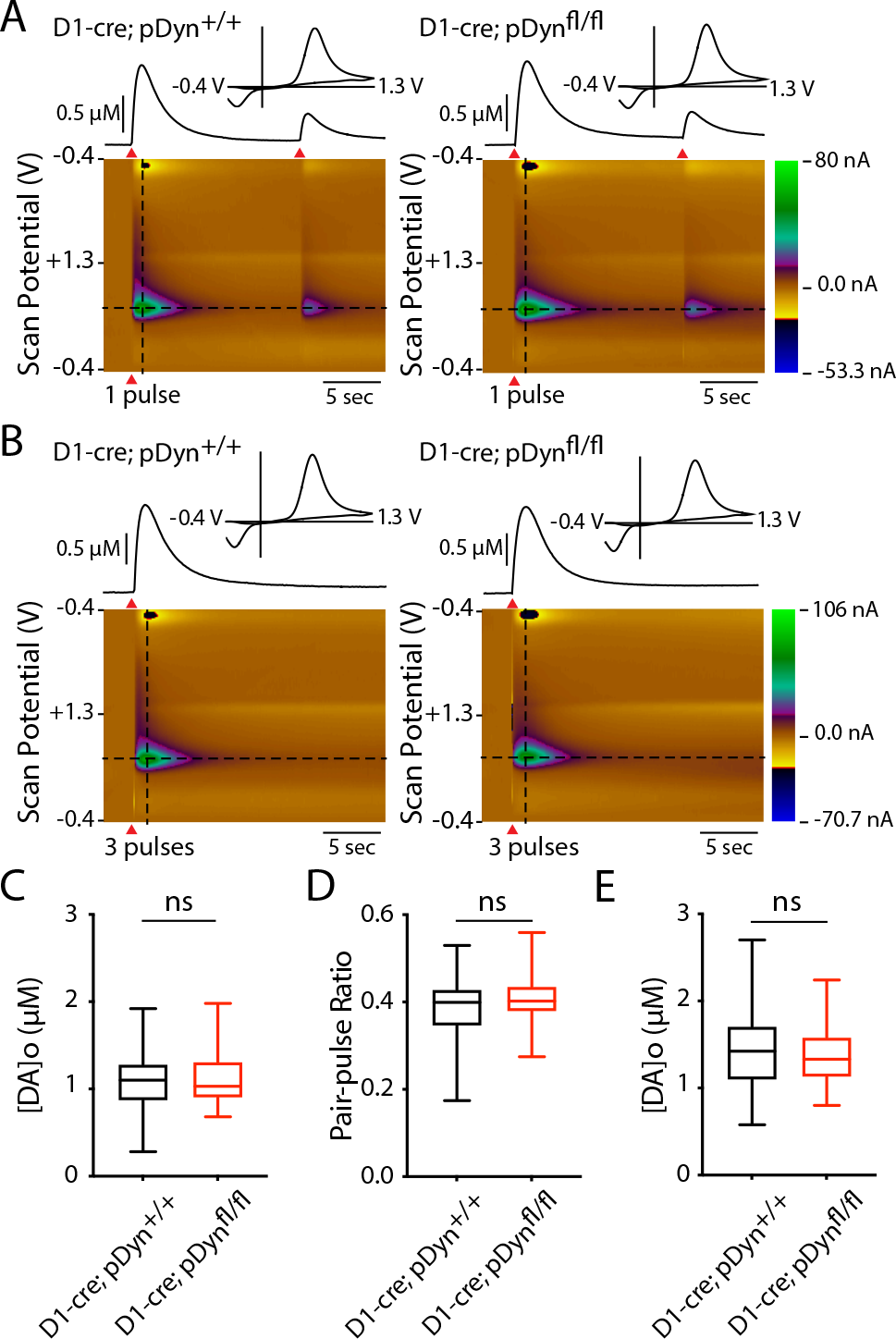
D1-pDyn cKO did not affect dopaminergic transmission. (**A, B**) Representative fast-scan cyclic voltammetric (FSCV) recordings of evoked DA response in a WT control and a cKO dorsal striatal slice. The recording traces show the responses evoked by two single pulses separated by 15s (A) or three 50 Hz pulses stimulation (B), collected at the horizontal dash line in the pseudo-color plot. Inset: traces showing the voltammogram profile of the responses, collected at the vertical dashed line in the pseudo-color plot. Red arrowheads indicate the stimulation timing. (**C**) Mean peak [DA]o evoked by 3 pulses are similar in WT and cKO groups. WT, n = 30 slices / 5 mice, 1.42 ± 0.081; cKO, n = 30 slices / 5 mice, 1.42 ± 0.073, p = 0.82, Mann-Whitney. (**D**) Left: mean peak [DA]o evoked by the first single pulse is not significantly different in WT and cKO slices. WT, 1.1 ± 0.061; cKO, 1.14 ± 0.066, p = 0.87, Mann-Whitney. Right: paired-pulse ratios of [DA]o evoked by two single pulses separated by 15s. WT, 0.39 ± 0.014; cKO, 0.41 ± 0.011, p = 0.43, Mann-Whitney. Data are presented as mean ± SEM. ns: not significant, p > 0.05.

**Figure S2.**
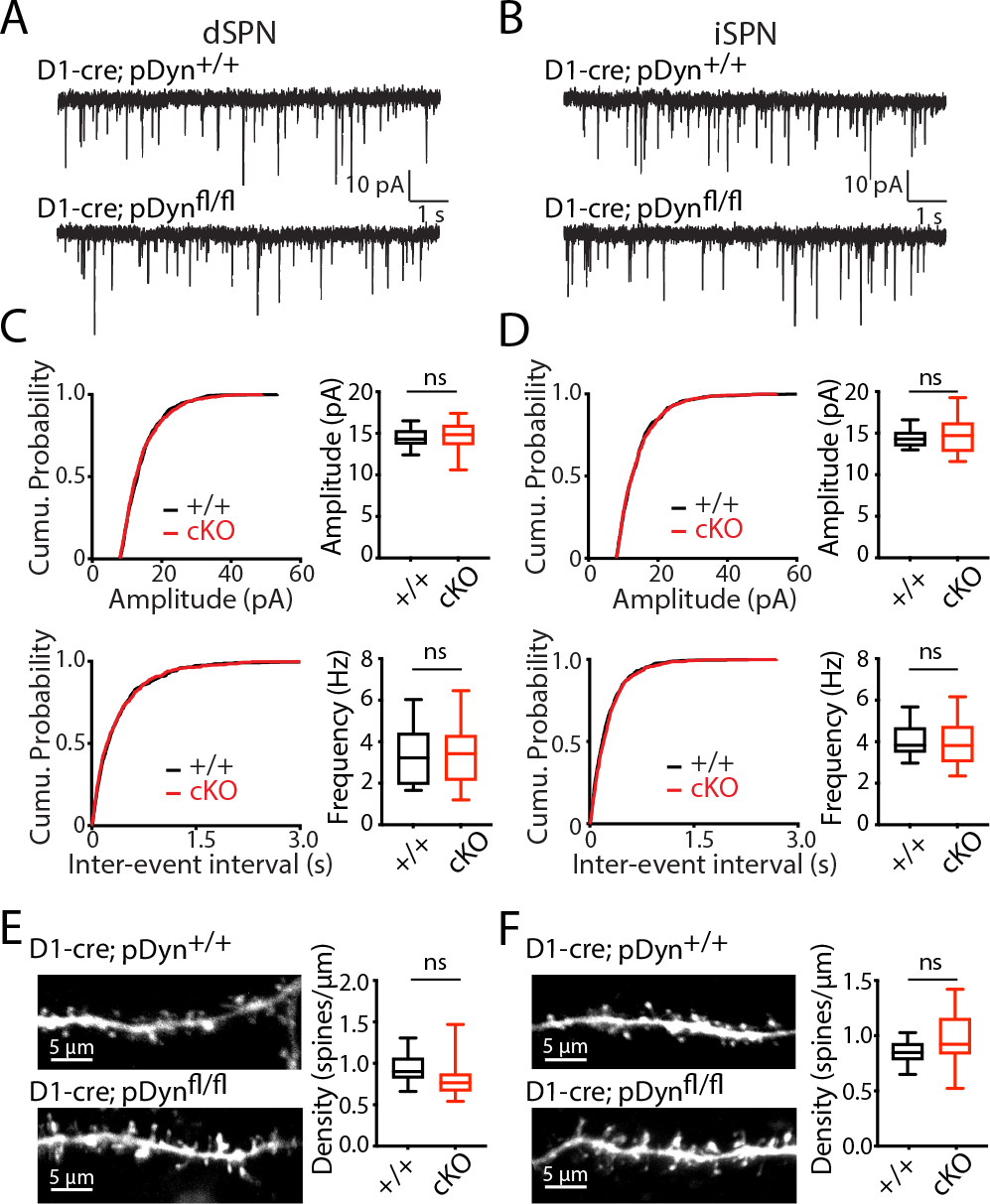
D1-pDyn cKO did not affect baseline glutamatergic transmission or spine density. (**A, B**) Representative mEPSC traces recorded from dSPNs (A) and iSPNs (B) from WT and cKO mice. (**C**) Cumulative probability distribution and bloxplot summary of mean EPSC amplitude (*upper*) and inter-event intervals (lower) recorded from dSPNs. There are no significant differences in dSPN mEPSC amplitude (left: p = 0.62, Kolmogorov-Smirnov test; right: mEPSCs amplitude: WT, 14.43 ± 0.32, n = 13/4; cKO, 14.68 ± 0.51, n = 13/2, p = 0.59, Mann-Whitney) or frequency between WT and cKO mice (left: p = 0.82, Kolmogorov-Smirnov test; right: WT, 3.27 ± 0.39, n = 13/4; cKO, 3.41 ± 0.43, n = 13/2, p = 0.87, Mann-Whitney). (**D**) Cumulative probability distribution and bloxplot summary of mean EPSC amplitudes (*upper*) and inter-event intervals (lower) recorded from iSPNs. There are no significant differences in iSPN mEPSC amplitude (left: p = 0.95, Kolmogorov-Smirnov test; right: mEPSC amplitude: WT, 14.33 ± 0.28, n = 14/4; cKO, 14.82 ± 0.61, n = 13/2; p = 0.78, Mann-Whitney) or frequency between WT and cKO mice (left: p = 0.33, Kolmogorov-Smirnov test; right: WT, 4.09 ± 0.24, n = 14/4; cKO, 3.99 ± 0.34, n = 13/2, p = 0.73, Mann-Whitney). (**E, F**) Left: Example images of spiny dendritic segments of dSPNs (E) or iSPNs (F) from WT and cKO mice. Right: Boxplot summary of spine densities of dSPNs (E) or iSPNs (F). There are no significant differences in dSPN and iSPN spine density between WT and cKO. dSPN: WT, n = 12/5, 0.93 ± 0.052; cKO, n = 11/4, 0.84 ± 0.083, p = 0.091, Mann-Whitney. iSPN: WT, n = 9/2, 0.8 ± 0.03; cKO, n = 14/4, 0.82 ± 0.052, p = 0.45 Mann-Whitney. Data are presented as mean ± SEM. ns: not significant, p > 0.05.

## Notes

### Competing Interest Statement

The authors have declared no competing interest.

